# ICGenealogy: Mapping the Function of Neuronal Ion Channels in Model and Experiment

**DOI:** 10.1101/058685

**Authors:** William F Podlaski, Alexander Seeholzer, Lukas N Groschner, Gero Miesenböck, Rajnish Ranjan, Tim P Vogels

**Affiliations:** Centre for Neural Circuits and Behaviour, University of Oxford, Oxford OX1 3SR, UK; Department of Physiology, Anatomy and Genetics, University of Oxford, Oxford OX1 3QX, UK; Laboratory of Computational Neuroscience, EPF Lausanne, 1015 Lausanne, Switzerland; The Blue Brain Project, EPF Lausanne, 1015 Lausanne, Switzerland

## Abstract

Ion channel models are the building blocks of computational neuron models. Their biological fidelity is therefore crucial for the interpretability of simulations. However, the number of published models, and the lack of standardization, make the comparison of models with one another and with experimental data difficult. Here, we present a framework for the automated large-scale classification of ion channel models. Using annotated metadata and model responses to a set of voltage-clamp protocols, we assigned 2378 models of voltage- and calcium-gated ion channels coded in *NEURON* to 211 clusters. The *IonChannelGenealogy* web interface provides an interactive resource for the categorization of new and existing models and experimental recordings. It enables quantitative comparisons of simulated and/or measured ion channel kinetics, and facilitates field-wide standardization of experimentally-constrained modeling.

## Introduction

Ion channels play crucial roles in neuronal signal processing (Koch and Segev, 2000; Cai et al., 2004; Goldberg et al., 2008) and plasticity (Sjostrom and Nelson, 2002; Shah et al., 2010; Debanne et al., 2003). Interactions among the many different ion channels expressed by a single cell can lead to extraordinarily complex dynamics, whose dissection necessitates computational modeling, as first demonstrated by Hodgkin and Huxley in 1952 for action potential generation (Hodgkin and Huxley, 1952). Simulation environments like *NEURON* (Hines and Carnevale, 2001; Carnevale and Hines, 2006) can be used to create biophysical models with realistic morphologies, ionic currents, and channel densities (Figure 1A), facilitating the integration of experimental data into models (Mainen and Sejnowski, 1995; Stuart and Spruston, 1998; Migliore et al., 1999; Poirazi et al., 2003; Destexhe and Pare, 1999; Traub et al., 2003). More than a thousand neuronal models, and several thousand individual ion channel models, are archived in the online database *ModelDB* (Hines et al., 2004), which enables other researchers to verify original claims, and to reuse and extend existing models in the light of new results.

**Figure 1.**
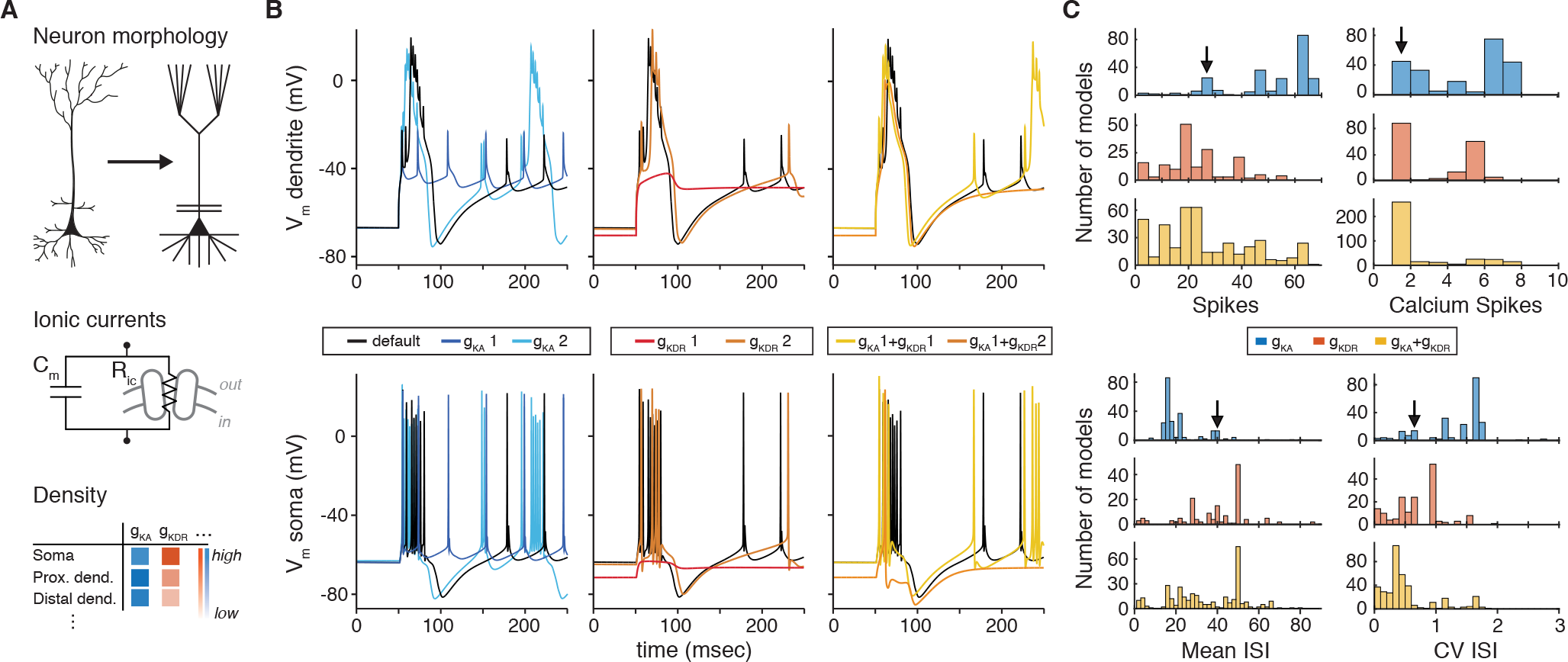
The choice of ion channel model influences the behavior of a simulated neuron. **A**: Biophysical neuron models are composed of a detailed multicompartmental morphology, several active ion channel conductances, and a density of each conductance that depends on the specific compartment. **B**: Simulation of a detailed layer 2/3 pyramidal neuron model, adapted from Traub et al. (2003) (see Methods for details). The model was stimulated with a 1.5nA current step beginning at 50msec, while recording the membrane potential in the apical dendrite (top) and soma (bottom). Simulations were first run using the original conductances from Traub et al. (“default”, black). Left: the default A-type potassium model (*g_KA_*) was replaced with two other A-type models (*g_KA_*1: dark blue, Hay et al. (2011), ModelDB no. 139653; *g_KA_*2: light blue, Traub (2004), ModelDB no. 45539). Middle: the default delayed rectifier potassium model (*g_KDR_*) was replaced with two other delayed rectifier models (*g_KDR_* 1: orange, Zhou and Hablitz (1996), ModelDB no. 3660; *g_KDR_*2: red, Durstewitz et al. (2000), ModelDB no. 82849). Right: both A-type and delayed rectifier models were replaced with other models (*g_KA_*1 + *g_KDR_* 1, yellow; *g_KA_* 1 + *g_KDR_*2, orange). **C**: Model from **B** was simulated for 1000msec with a 1.5nA current step, replacing the default A-type current with each A-type-labeled model on ModelDB (243 total models, blue), then replacing the default delayed rectified current with each delayed rectifier model on ModelDB (188 total models, red), and finally replacing both A-type and delayed rectifier default currents with a sample of combinations of others (441 combinations, yellow). Summary measures are shown for total number of spikes, total number of calcium spikes, mean inter-spike interval (ISI) and coefficient of variation (CV) of ISI during the 1000msec period. Black arrows represent the simulation results for the default model.

Matching model and experiment is essential for biophysical models, in which many components have a direct biological counterpart (Brette et al., 2007). For example, pyramidal neuron models have been shown to reproduce the recorded spiking activity of these cells accurately with a particular set of ion channels (Traub et al. (2003); Figure 1B, black traces; see Methods). However, the dynamics can change, sometimes dramatically, when one of the modeled ion channel currents is exchanged for an identically-labeled model from a different publication on *ModelDB* (Figure 1B, colored traces, Figure 1C). This example underscores the importance of selecting ion channel models, yet there is currently no standardized experimental dataset against which to validate them.

Furthermore, the increasingly large number of models on ModelDB (e.g., over 300 new ion channel models in 2014 alone; Shepherd Lab (2015)), with non-standardized labeling and a high degree of redundancy, makes it difficult to understand how models relate to each other and to biology. For example, a researcher looking to use an existing *A-type* potassium channel model will find over 250 *A-type* models, spanning a range of behaviors (Figure 1C, blue). Instead of a thorough and time-consuming fitting of appropriate ion channel dynamics, it is common for modelers to adapt previously published models for their own purposes. However, this may introduce experimentally unverified systematic changes or even errors into later generations of models and may have dramatic effects on the biological interpretability of the results.

To facilitate informed choices among this bewildering variety of models, we categorized 2378 published voltage- and calcium-dependent ion channel models in *NEURON* that are available on *ModelDB*. We cataloged all relevant information about each ion channel model from the associated literature, including its *pedigree* relations: whether a given model is based on previous models, and, if so, which ones. Additionally, we compared the kinetics of each ion channel model in standardized voltage-clamp protocols. The resulting maps of ion channel behavior show model variability and diversity, and point to the computational and experimental sources that were used to fit each model. Our efforts have grouped 2378 ion channel models into 211 clusters, dramatically simplifying the search for an appropriate ion channel model.

We present our findings in an annotated, interactive web-interface with a short video manual (ICGenealogy (2015); http://icg.neurotheory.ox.ac.uk), that allows filtered search of individual ion channel models by metadata and relational information, and the comparison of channel model kinetics. The underlying database is freely accessible via a web API. In an effort to make our resource compatible with experimental data and new models, we offer the possibility to upload and assess the similarity of experimentally recorded current traces (as well as new models and model traces) in the same topology. We show an example of the use of this comparison through the analysis of an unclassified ion channel model, as well as an experimentally recorded voltage-dependent potassium current from *Drosophila melanogaster*. In summary, we provide a framework for the direct and automated comparison of models and experiments to facilitate experimentally constrained modeling and quantitative characterization of ion channel behavior.

## Results

### Categorizing ion channel models by metadata and ancestor-descendant relationships

To build a map of ion channel model function, we categorized and analyzed a widely-used subset of 2378 voltage- and calcium-dependent ion channel models (*".mod"* files) in the *NEURON* simulation environment (Hines and Carnevale, 2001; Carnevale and Hines, 2006). A set of “metadata” was extracted manually from the associated journal articles for each model file (Figure 2A, top): reference information (Ref. Info, including author(s) of the model code), ion channel information (I.C. info: ion selectivity, gating mode, subtype), system information (Sys. Info: brain area, neuron type, neuron region, animal model), as well as additional comments (Other: e.g. temperature constraints).

**Figure 2.**
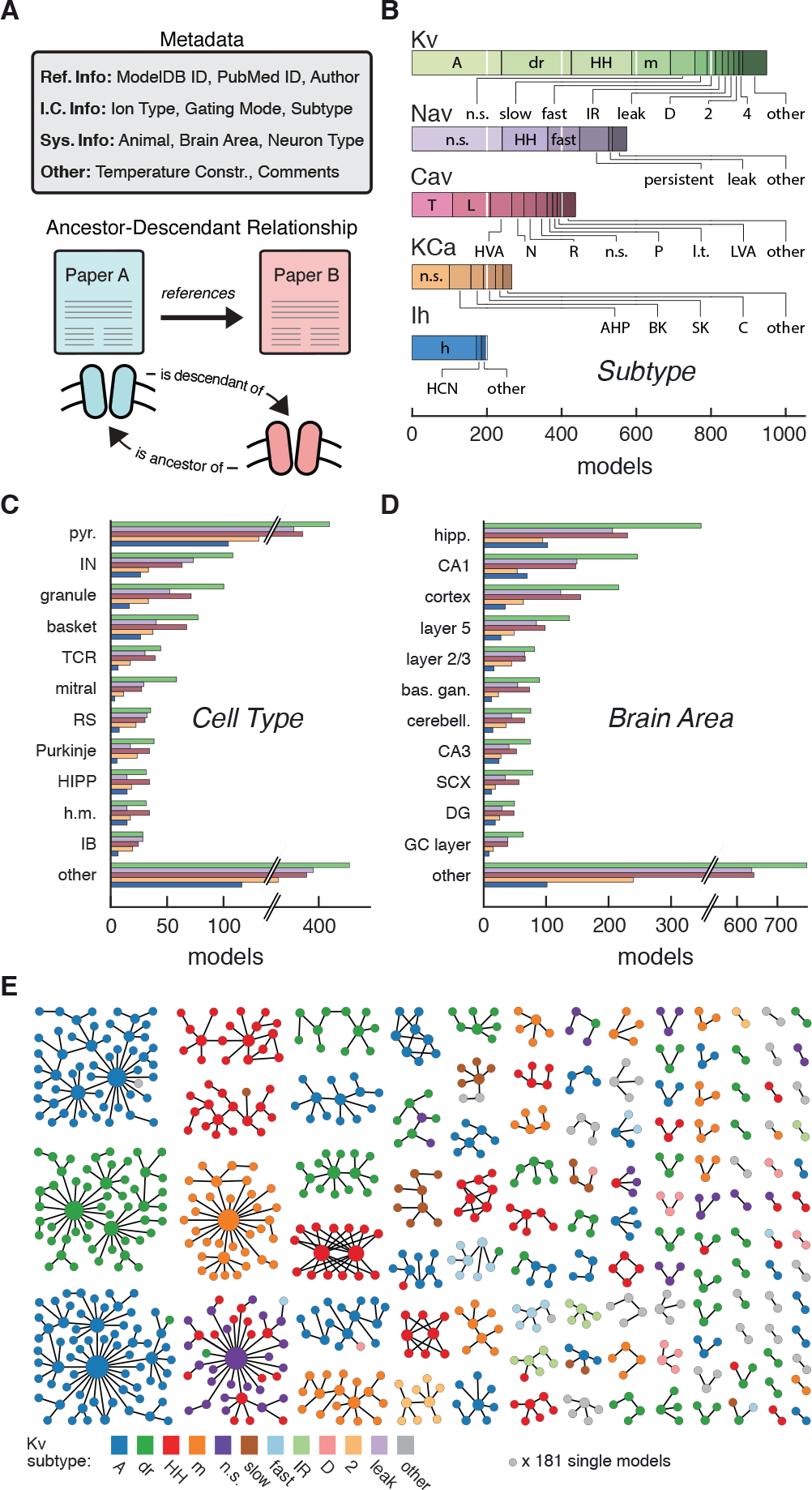
Ion channel models can be categorized by metadata and ancestor-descendant relationships. **A**: Metadata were manually extracted from ModelDB and associated journal articles (top). Ancestor-descendant relationship (bottom) was established between different models (see main text for description). **B**: Models were divided into five classes based on ion type: voltage-dependent potassium (Kv), voltage-dependent sodium (Nav), voltage-dependent calcium (Cav), calcium-dependent potassium (KCa), and hyperpolarization-activated cation (Ih). Each class is divided into subtypes, ordered from left to right according to group size. Uncommon subtypes are grouped together (other). **C**, **D**: Histogram of cell types and brain areas for each ion type, ordered from top to bottom by the number of models. Colors as in **B**. **E**: Pedigree graph displaying families of the Kv class, sorted by family size. Each node represents one model, colored by subtype, and edges represent ancestral relations between models (panel A, bottom). Note that unconnected models (181 total) are not shown. A: A-type, dr: delayed rectifier, HH: Hodgkin-Huxley, m: m-type, n.s.: not specified, IR: inward rectifier, HVA: high-voltage activating, N: N-type, R: R-type, P: P-type, l.t.: low threshold, LVA: low-voltage activating, AHP: after-hyperpolarization, BK: big conductance, SK: small conductance, HCN: Hyperpolarization-activated cyclic nucleotide-gated, pyr.: pyramidal, IN: interneuron, TCR: thalamocortical relay, RS: regular spiking, HIPP: hilar perforant-path associated, h.m.: hilar mossy, IB: intrinsic bursting, hipp: hippocampus, bas. gan.: basal ganglia, cerebell.: cerebellum, SCX: somatosensory cortex, DG: dentate gyrus, GC: granule cell.

Based on ion selectivity and gating mode, the majority of all ion channel models (~75%) fell into five classes (Figure 2B): voltage-dependent potassium (Kv), voltage-dependent sodium (Nav), voltage-dependent calcium (Cav), calcium-dependent potassium (KCa) and hyperpolarization-activated cation (Ih) channel models. We recorded 74 different subtype labels across all classes (Kv: 32, Nav: 19, Cav: 20, KCa: 11, Ih: 5; Figure 2B, cf. Table S1). Prominent modeled neuron types were *pyramidal*, *interneuron*, *granule cell*, and *basket cell* (Figure 2C), and prominent brain areas included *hippocampus*, and *cortex* (Figure 2D). Other metadata also showed a broad variety across channel models (ICGenealogy, 2015).

To denote family relations (Figure 2A, bottom), model *A* was labeled as a “descendant” of an older “ancestor” model *B* if the publication reporting *A* cited the publication for *B* as the source or inspiration of its channel dynamics, or if the code of Models *A* and *B* were sufficiently similar (Methods). Visualizing family relationships makes it apparent that many models form large families, often with a highly-cited hub model that has many descendants (Figure 2E). On the other hand, there are a large number of small families and model singletons that imply *de novo* model creation, lack of appropriate citations, or translation from other simulators (noted in the metadatum *comments*). Subtype labeling mapped well onto families, but family identity did not guarantee homogeneity of subtype or vice versa - all individual subtypes were found across several families (see Figure 2E for Kv, and Figures S1–Figures S2A,Figures S2F for other ion type classes).

Family relationships and metadata thus help to distinguish ion channel models, but the lack of standardized annotations in a common nomenclature, as well as the sheer abundance of models make it difficult to infer the degree of their functional diversity. Based on metadata alone, it is thus difficult to choose a model for appropriation into one’s own work.

### Defining functional groups of models through voltage clamp protocols and clustering

To quantify the functional relationships between channel models, we used a set of voltage-clamp simulation protocols, in kind with those developed for the experimental characterization of ion channels (Sakmann and Neher, 2009; Ranjan et al., 2011). We chose this procedure to assess the spectrum of possible dynamics in a model-free manner, i.e. without explicitly taking into account the underlying equations. This allows for the comparison of models strictly based on their behavior, and, as we discuss later, the direct comparison with experimental data.

Using the *NEURON* simulation environment (Hines and Carnevale, 2001; Carnevale and Hines, 2006), each ion channel model was placed individually into a model soma and its current responses to five voltage-clamp protocols were recorded (Figure 3A, left; see Methods, Figure S3, Table S2 for full description). The protocols were designed to probe the gating characteristics of ion channels, i.e. activation, deactivation and inactivation, as well as temporal dynamics during voltage ramping and repeated action potentials. Protocol parameters were adjusted for each of the five ion classes separately. Current responses were normalized to remove the dependence on the maximum conductance, and subsampled at particular regions of interest (Figure 3A, dashed areas) to obtain a trace of characteristic data points for each protocol (Figure 3B, Figure S3F). Using principal component analysis (PCA) across all traces of a particular ion channel type, we obtained a final *D*-dimensional score for each model, accounting for at least 99% of the variance across all channel models in each class. The dimensionality *D* varied between 16 and 29 dimensions for the five classes (Kv: 16, Nav: 21, Cav: 29, KCa: 16, Ih: 16). The Euclidean distance between any two given model scores was termed their “similarity” (Figure 3C, top). Finally, we used Ward’s clustering method (Ward, 1963) on the model scores to establish an agglomerative hierarchy of ion channel model clusters (Figure 3C, bottom).

**Figure 3.**
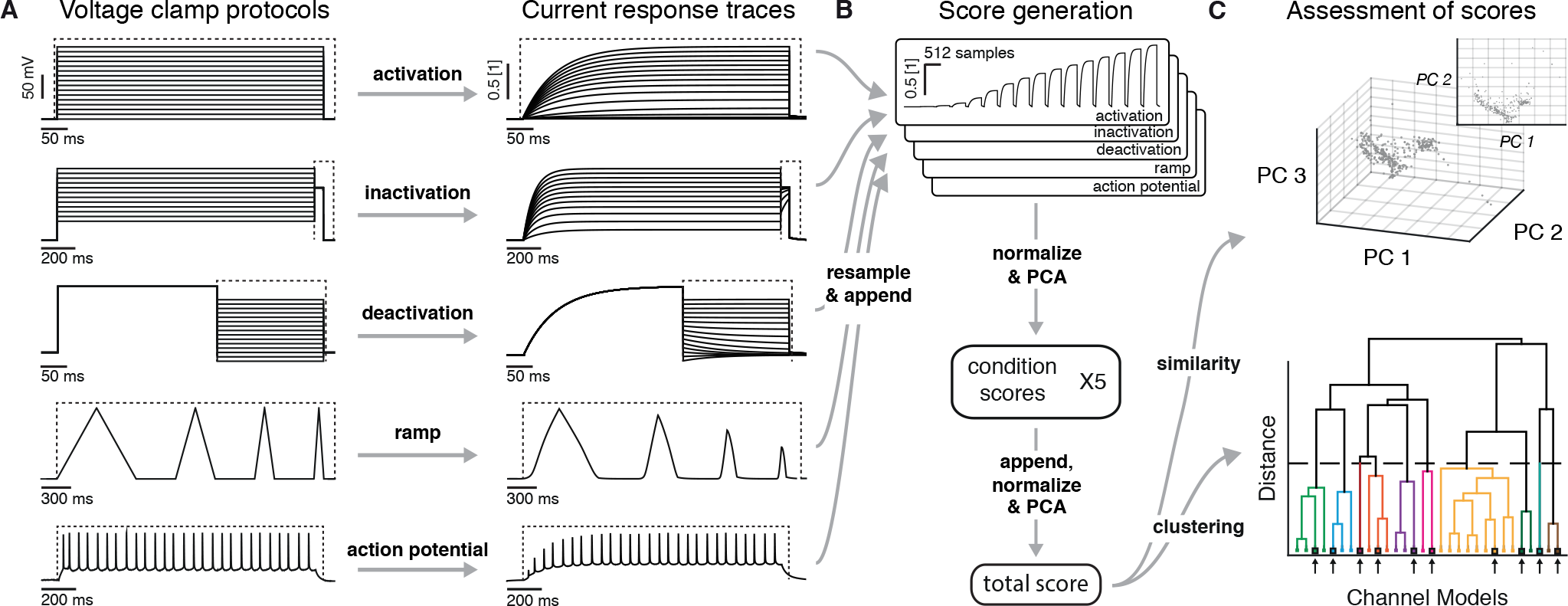
Voltage-clamp protocols for the quantitative analysis of ion channel models. **A**: Left: five voltage clamp protocols were used to characterize ion channel responses recorded in single compartment somata simulated in *NEURON* (see Figure S3 for full description). Multiple lines indicate a series of increasing voltage steps with the same time sequence. Right: current response traces are shown for an example model. Dashed regions indicate response times used for data analysis. **B**: Current responses were subsampled and appended, then dimensionality-reduced by principal component analysis (PCA) to form a *condition score* vector for each protocol. These score vectors were further normalized and dimensionality-reduced to form a *total score* vector. **C**: The first three principal components of the score vector are shown for Kv ion channel models (top). Scores were clustered using an agglomerative hierarchical clustering technique (bottom). Distinct clusters (noted by colors) form when a cutoff (dashed line) is introduced in the distance between hierarchical groupings, chosen based on several cluster indexes (see Figures S4–Figures S5). *Cluster representative models* (bold squares with arrows) are selected as reference models for each cluster (see Methods).

An appropriate number of clusters was obtained through a variety of published cluster indexes (Methods, Figures S4–Figure S5). For the Kv class, this resulted in 60 clusters with distinct responses (Figure 4) and small intra-cluster variability (Figure S6). The other classes divided similarly into 38, 43, 44, and 26 clusters for Nav, Cav, KCa, and Ih, respectively (see Figures S1–Figure S2). We named clusters according to the most common label of their members and we denoted the model closest to the mean score coordinate of each cluster as its reference model.

**Figure 4.**
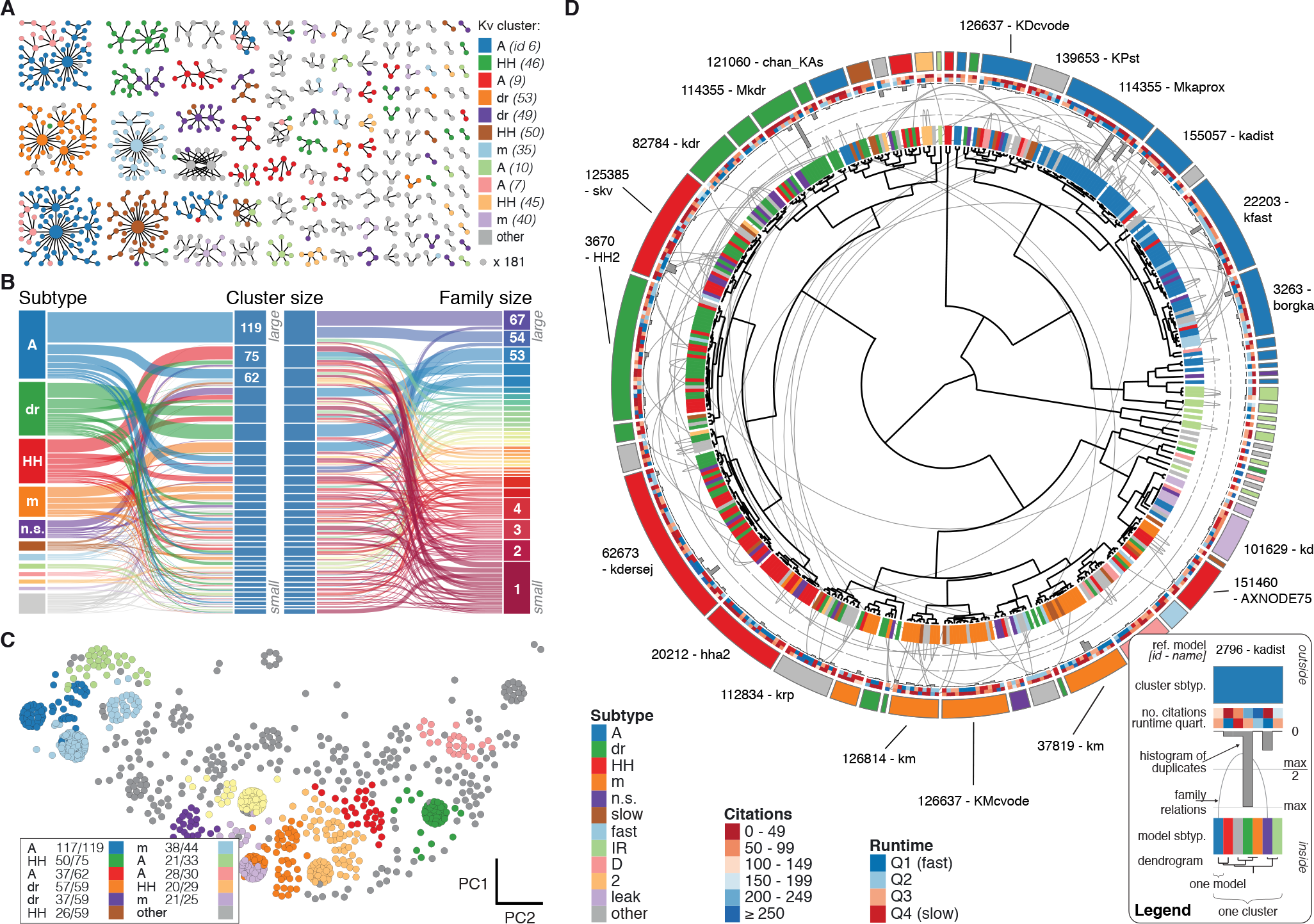
Quantitative analysis of Kv ion channel models: functional map and clusters of common behavior. **A** Pedigree graph of the Kv class (cf. Figure 2E), colored by membership in the 11 largest clusters in the class (named by most prevalent subtype, bottom). Membership to other clusters is indicated by grey color. Cluster ID is given in parentheses for easy comparison with website. **B**: ‘Sankey’ diagram for the Kv ion type class, showing the relation between subtype, cluster identification and family identification, each ordered from top to bottom by increasing group size. The 11 most common subtypes are shown in color, with all others grouped together in grey. Small families (size 1 to 6 members) are grouped together. **C**: Plot of Kv models in the first two principal components of score space. Colors indicate membership in one of the 11 largest clusters in the class, with membership to other clusters colored in grey. Clusters are named by their most common subtype, with the proportion of that subtype specified in the legend. Points lying very close to each other have been distributed around the original coordinate for visualization. D: ‘Circos’ diagram of the Kv ion type class. All unique ion channel models are displayed on a ring, organized by cluster identification. From outside to inside, each segment specifies: cluster reference model (only displayed for large clusters), cluster subtype, number of citations, runtime, number of duplicates, model subtype, as well as a dendrogram of cluster connections (black) and family relations (grey). A: A-type, dr: delayed rectifier, HH: Hodgkin-Huxley, m: m-type, n.s.: not specified, IR: inward rectifier. See Figures S1 and S2 for other ion type classes.

We found that most ancestor-descendant families fell within one cluster, indicating consistency between the family relations collected from the papers and model behavior (Figure 4A–Figure 4B, Figures S1–Figure S2B,Figure S2C and Figure S2G,Figure S2H). However, a common subtype label did not guarantee a common cluster identity (Figure 4B, Figures S1–Figure S2C,Figure S2H). Many models with the same subtype fell into different clusters. For example, the ~250 A-type-labeled Kv ion channel models fell into 14 clusters (although only five clusters comprised over 90% of them, Figure 4B). These clusters contained few other subtype labels, suggesting that *A-type* is generally a consistent label for at least five similar, yet distinct kinetic behaviors. Moreover, the similarity between these clusters was generally high (and thus they were plotted within the same vicinity on the wheel of the ‘Circos’ plot, Figure 4D; see also Methods). Other subtype labels across all ion channel types showed similar results (Figures S1–Figure S2C,Figure S2H). Interestingly, for four of the five ion type classes (KCa being the exception), most isolated single-model clusters corresponded to genealogical singletons, supporting the idea that these models are indeed unique, and do not appear isolated simply due to missing ancestor-descendant links. However, this was not true for all genealogical singletons, many of which were kinetically aligned to larger families and consequently fell into the same clusters. In conclusion, clustering allowed us to identify 211 distinct groups of ion channel models that share similar behavior, regardless of publication context or subtype labeling.

### Ion channel model groups defined by common metadata show variability in behavior

The variability in behavior of identically-labeled models in different clusters may stem from various sources. Biological diversity and differences in experimental setup would explain why ion channels of the same subtype have been fitted by different models with varying kinetics. Consistent with this notion, the behavior of groups of models defined by common subtype, neuron type and brain area (Figure 5A, plotted data points) is often more diverse than that of models defined by a common cluster (Figure 5A, dashed line).

**Figure 5.**
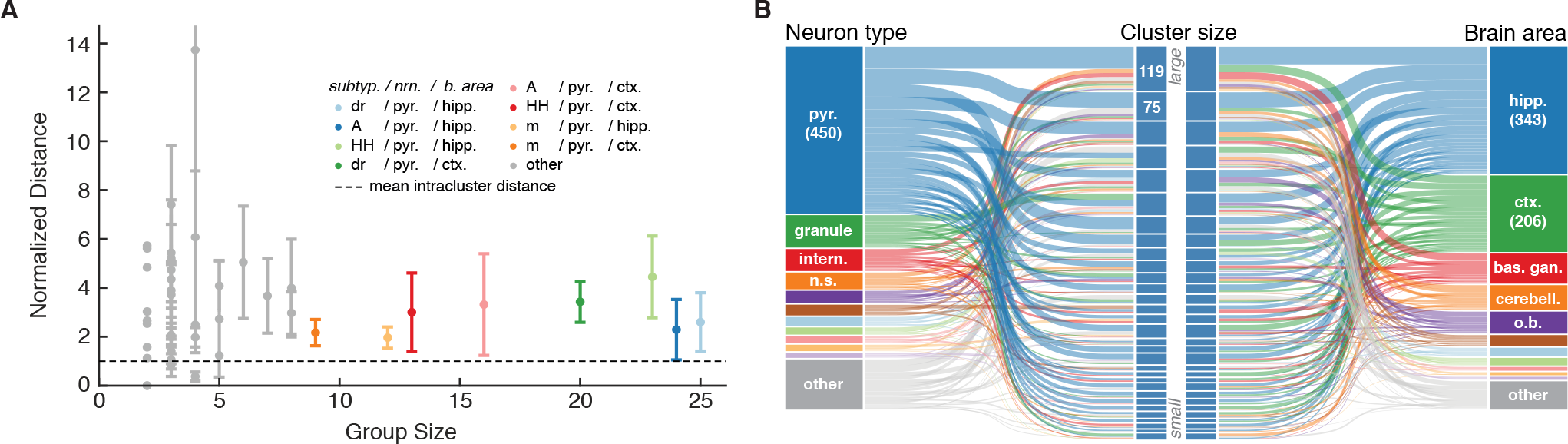
Ion channel model groups defined by common subtype, neuron type and brain area show variability in behavior. **A**: Kv models are grouped by common subtype, neuron type, and brain area. The mean pairwise distance in score space between all models within a group is plotted against group size, with errorbars corresponding to the first and third quartiles (middle 50% of distances). The largest eight groups are shown in color. Distances are normalized relative to the mean pairwise distance between models in the same *cluster* (dashed line; averaged across all clusters with at least one pairwise distance greater than zero, 43 of 60 clusters). Most groups have a larger mean distance than the average cluster (above the dashed line). **B**: ‘Sankey’ relational diagram for metadata, neuron type, cluster identification and brain area, for the Kv ion type class. Bars are stacked histograms, i.e. the height of bars indicate the relative number of models. Left column: partition of channels by 11 most prevalent neuron types. Middle column: partition of channels into assigned clusters. Right column: partition of channels by 11 most prevalent brain areas. Links between columns are colored according to neuron type (left) and brain area (right). Numbers in left/right column are number of channels per label. pyr: pyramidal, intern.: interneuron, n.s.: not specified, hipp.: hippocampus, ctx: cortex, bas. gan.: basal ganglia, cerebell: cerebellum, o.b.: olfactory bulb.

A large portion of the variability, however, may stem from model fitting and idiosyncratic changes to individual model implementations. We find no clear correspondence of any given cluster with categories such as brain area and neuron type (Figure 5B). Nearly every cluster contains models that have been used in pyramidal cells (Figure 5B, left, blue) of both cortex and hippocampus (Figure 5B, right, blue & green). This suggests that the variability in models does not reflect the biological diversity of ion channels across different biological systems. In the same vein, e.g., *A-type*-labeled models that have been used in pyramidal cells of the hippocampus (117 models) are found in 9 clusters (cf. ICGenealogy (2015)). Finally, references to experimental data used to fit models can also be found in multiple clusters.

### Automated comparison of new models and experimental data with the database

Our database and analysis framework, accessible through the web interface, enable the automated analysis of new computational models as well as experimental data (Figure 6A). To illustrate this process, we uploaded and tested a previously uncatalogued Kv model from a hippocampus CA1 pyramidal cell model (*kad.mod* from Hsu et al. (2015); ModelDB ID no. 184054). We compared its scores and response traces to the presently available 931 Kv models (Figure 6B) and determined its relation to previous ion channel implementations. We found that the model fits well within a cluster of mostly pyramidal A-type-labeled models used in simulations of rodent hippocampus, thereby verifying the assumed characteristics of the model.

**Figure 6.**
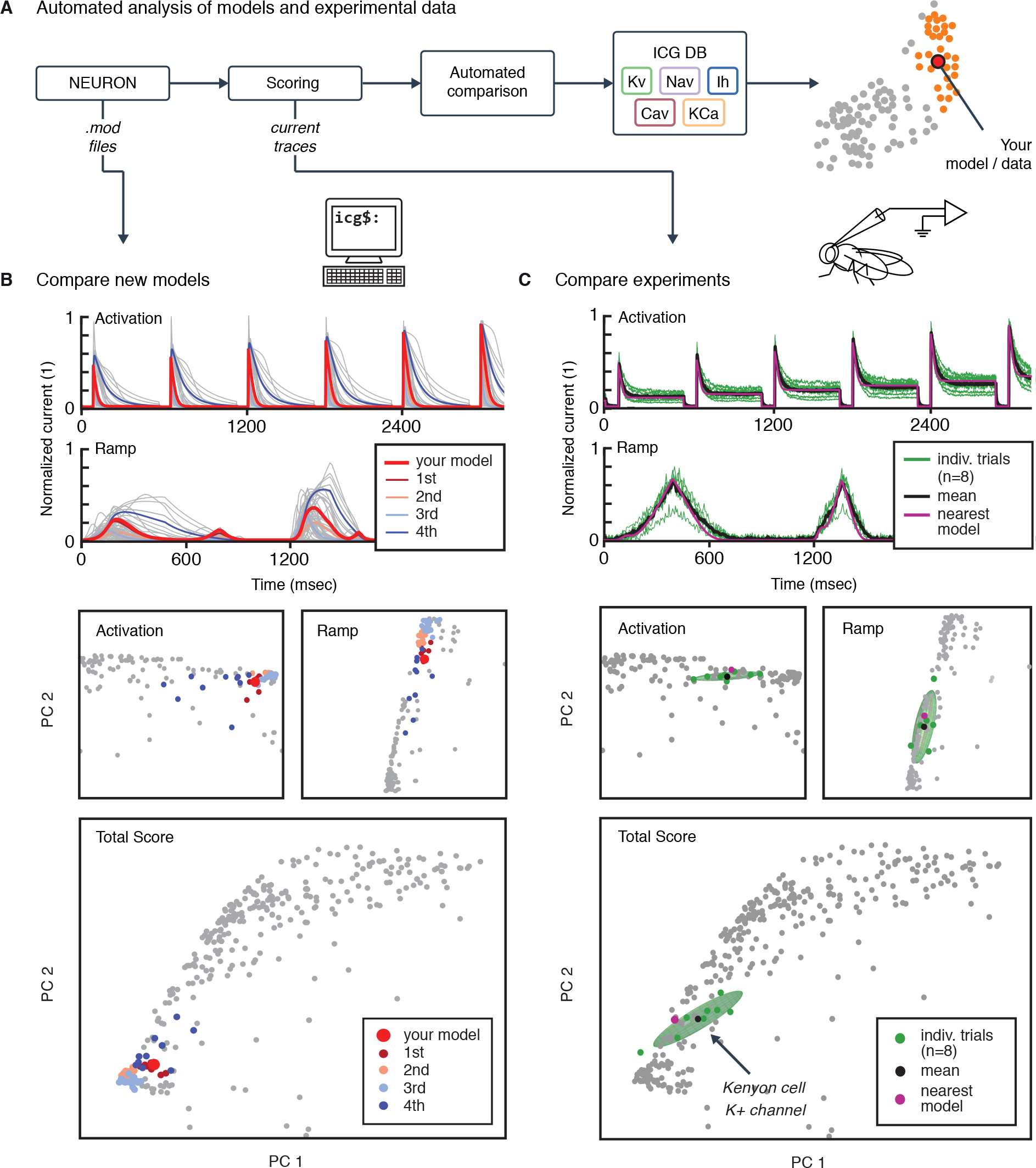
Automated analysis of new models and experimental data. **A**: Flowchart of data processing steps involved in automated comparison. Source code for model files written in *NEURON* can be uploaded to the website, and current responses are automatically generated. Current traces are processed to compute scores, which are compared to all models in the database (illustrated in **B**). Additionally, raw current traces obtained experimentally (or from models in other languages) can be uploaded and analyzed directly (illustrated in C). **B**: Example analysis and comparison of a new ion channel model *(kad.mod* from Hsu et al. (2015); ModelDB ID no. 184054). Top: Segments of the current response traces (red) for activation (voltage steps 10 - 60mV) and ramp protocols (first half), along with the closest four clusters (other colors: mean currents, grey lines: individual currents). Bottom: first two principal components of score space for activation and ramp protocols, as well as total score. **C**: Example analysis of in vivo recordings of a K+ current from Drosophila Kenyon cells (see Methods for details and Figure S7 for full traces) and comparison to ICG database. Top: mean (n=8 recordings, black) and individual recordings (green). Bottom: mean (black dot) and individual experimental recordings (green dots) plotted in the first two principal components of score space (ellipsoid illustrates the variance across individual recordings). Comparison is made to the nearest (in score space) ion channel model in the database (magenta; *Kv4_csi*, ModelDB i.d. 145672). Grey dots in **B** and **C** are scores of Kv channel models in the database.

The framework can also be used for the comparison of experimental data and models. To illustrate, we uploaded and tested an experimental dataset of recordings from Kenyon cells in *Drosophila melanogaster*. Voltage-gated cationic currents across the membranes of these neurons are thought to be dominated by A-type K+ channels, in particular Shal/Kv4 (Gasque et al., 2005). This renders Kenyon cells a suitable neuronal cell type to test the biological relevance of our voltage-clamp protocols in an *in vivo* setting. Current responses were recorded in targeted whole-cell patch clamp experiments *in vivo* (see Methods). The recordings were performed using our five standardized voltage-clamp protocols, allowing us to transform and compare the experiments directly to models in the same ‘score’ space (Figure 6C). The comparison revealed a close match to an existing model in the database, and thus characterizes the behavior of the ion channel as similar to a mammalian Kv4 ion channel (Figure 6C; Fineberg et al. (2012); *Kv4_csi*, ModelDB i.d. 145672).

## Discussion

Neuroinformatics has become an increasingly important part of neuroscience research, as new technology and large-scale research projects push the field into the realm of *big data* (Akil et al., 2011; Ferguson et al., 2014; Grillner, 2014; Tripathy et al., 2014). Importantly, the need for assessment and aggregation of published knowledge extends beyond experimental data, and has recently started encompassing computational models of neural function (Hines et al., 2004; Gleeson et al., 2013). Here, we have performed a meta-analysis of voltage– and calcium-dependent ion channel models coded in the *NEURON* programming language available in the database *ModelDB* (Hines et al., 2004). Our approach of combining metadata extracted from publications with a kinetics-based analysis allowed us to provide detailed information regarding the identity of each model in the database, filling in missing or ambiguous data, and validating the functional properties of channel models against their sometimes ambiguous nomenclature. Furthermore, we provide a framework for the large-scale comparison of models, both with each other and with experiments using the same standardized protocols, thus paving the way towards a unified characterization of ion channel function.

The voltage-clamp protocols used in this study were designed to efficiently probe the kinetics of all ion channel types considered here (Sakmann and Neher, 2009; Burgess et al., 2002; Hering et al., 2004). Measure the kinetic responses of each channel model allowed us to compare models regardless of their specific implementation. Notably, our method is amenable to the addition of other protocols that may be better suited to separate certain models.

Additionally, our analysis can be extended beyond the selection of channel models considered here. We limited our analysis to voltage-dependent and calcium-dependent ion channel models coded in the *NEURON* language, but, given the appropriate protocols, other types of ion-channels can be included. The same protocols can also be used to integrate models written for other simulators, or even simulator-independent formats, e.g., NeuroML (Cannon et al., 2014). We have taken steps to integrate our database and visualizations tightly with existing online resources, notably *ModelDB* (Hines et al., 2004).

The end result of our work is a dramatically reduced group of candidate models to test when looking for particular ion channel dynamics. Of the 2378 models in our database, we could identify 1132 models as unique, and further reduced this to 211 groups with substantially different kinetics. However, this does not eliminate the task of finding the most appropriate model, and we stress that the partition of channel models into clusters of similar response properties does not imply that models in the same cluster are necessarily redundant. Intra-cluster differences may still be important depending on the particular simulation at hand. Since the responses of different channels vary slowly and continuously rather than in discrete steps along the dimensions of the manifold of scores (see e.g. Figure 4B), the data may also be amenable to more sophisticated clustering and machine-learning approaches. To this end, the raw response data and scores have been made publicly available (ICGenealogy, 2015).

We provide an interactive database browser (ICGenealogy, 2015), which acts as a complement to existing resources such as ModelDB. It allows the comparison of channel models in five views: a similarity view focusing on the channel’s response kinetic scores (Figure 7A), a hierarchical tree view focusing on genealogical data (Figure 7B, top), an XY view to sort data by a given set of metadata dimensions (Figure 7B, middle), and a circular cluster view (Figure 7B, bottom). All these views feed a central comparison tool (Figure 7C), in which the metadata and traces for user-selected channel models can be viewed side-by-side. For specific examples of how to utilize this browser and to search for specific ion channel models, please refer to the instruction video (ICGenealogy, 2015) and manual (SOM).

**Figure 7.**
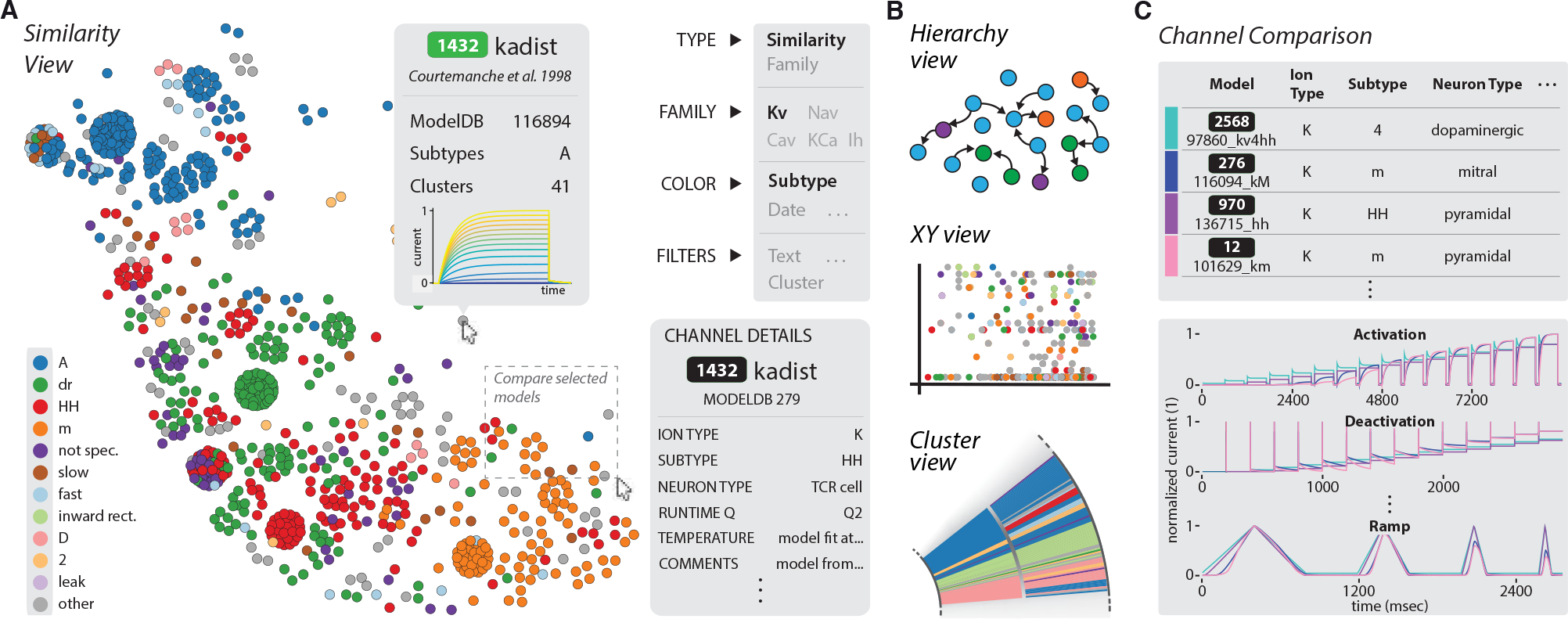
The ICGenealogy website allows for the interactive visualization of all data and analysis on the database (ICGenealogy, 2015) (http://icg.neurotheory.ox.ac.uk). **A**: Schematic of similarity view on website. Channel models of the Kv family are displayed in the first two principal components of score space, colored by subtype (legend on left). Hovering over models brings up information tooltip (center), and clicking on a model displays Channel Details (bottom right). Selected models can be compared by click-and-drag (see instruction manual SOM for more details). **B**: Schematic of other three views available on the website. Hierarchy view: models are displayed in a graph with edges that represent family relations. XY view: any two selected metadata are plotted against each other. Cluster view: models are organized in a ring partitioned by clusters. **C**: The channel comparison displays selected channels side-by-side with metadata (top) and current responses (bottom).

Because our voltage-clamp protocols are inspired by experimental procedures, models can be compared directly to experiments in an automated fashion. We have taken the firsts steps in this direction by showing a comparison of both a new model and an experimental dataset to the database here (Figure 6). While it is beyond the scope of the current study to integrate ion channel information from IUPAR/BPS Guide to Pharmacology (Pawson et al., 2014), Channelpedia (Ranjan et al., 2011) or other sources, these are important future steps which would help standardize nomenclature. Beyond its usefulness for cataloguing ion channel model behavior and pedigree, our resource will enable better experimentally-constrained modeling, and presents a first step towards a unified functional map of ion channel dynamics in model and experiment.

## Acknowledgements

The channel browser web-frontend was created by Phyramid (www.phyramid.com). We thank V. Harbuz and P. Toader from Phyramid for helpful discussions about visualization. We also thank P. Gleeson, S. Crook, J. Kwag, N.A. Cayco-Gajic and Y. Poirazi for helpful comments. Research was supported by grants from the Wellcome Trust Research Fellowship (G.M. and T.P.V.). A.S. was supported by the Swiss National Science Foundation (200020_147200). R.R. was supported by the EPFL Blue Brain Project Fund and ETH Board funding to the Blue Brain Project. L.N.G. was supported by a Wellcome Trust OXION studentship.

### Author Contributions

W.F.P. and A.S. contributed equally to this work. R.R. and T.P.V. conceived the original idea. W.F.P. and A.S. collected the data. W.F.P., A.S., R.R., and T.P.V. developed and performed analysis and visualization. A.S. created the database, backend and website. L.N.G. and G.M. provided experimental data. All authors wrote the manuscript.

## Experimental Procedures

### Pyramidal cell model (Figure 1)

A model of a layer 2/3 pyramidal cell was adapted from a previous study (Traub et al., 2003), ModelDB accession no. 20756. It contained 68 soma-dendritic compartments and 6 axonal compartments, with the following active conductances: leak (*g_L_*), transient (inactivating) Na+ (*g_NaF_*), persistent (noninactivating) Na+ (*g_NaP_*), delayed rectifier K+ (*g_KDR_*), transient inactivating K+ (*g_KA_*), slowly activating and inactivating K+ (*g_K2_*), muscarinic receptor-suppressed K+ (*g_KM_*), fast voltage– and calcium-dependent K+ (*g_KC_*), a slow calcium-dependent K+ (*g_KAHP_*), low-threshold inactivating Ca2+ (*g_CaT_*), high-threshold non-inactivating Ca2+ (*g_CaH_*), hyperpolarization-activating cation conductance (*g_Ih_*). We refer the reader to Traub et al. (2003) for channel kinetics, distribution and other details of the model.

The neuron model was simulated in the *NEURON* simulation environment (Hines and Carnevale, 2001; Carnevale and Hines, 2006), with a current step input injected into the apical dendrite, following Figure 2 of Traub et al. (2003). The protocol was as follows: 400 msec at −0.15 nA, followed by 1000 msec at 1.5 nA. A subset of this trace is shown in Figure 1B, comprising 50msec at −0.15nA and the first 200 msec at 1.5nA. The black traces in Figure 1B show the default behavior of the neuron model in response to injected input. The following four measures were computed for each spike train: total number of spikes, total number of calcium spikes, mean inter-spike interval (ISI) and coefficient of variation of ISI.

The stimulation paradigm was repeated in the presence of alternate ion channel models for *g_KA_*, taken from ModelDB (243 models total). All other ionic conductances, parameters and distributions remained the same. This was further done with alternate ion channel models for *g_KDR_* in a separate simulation (188 models total), with *g_KA_* set back to the original model. Finally, this was done in the case of replacing both *g_KA_* and *g_KDR_* currents. A subset of 441 pairs of alternate models were run together.

### The *ModelDB* database, *NEURON* language and the database nomenclature

The *ModelDB* database archives published neuron and network models (Hines et al., 2004). It contains over 1000 entries, with thousands of ion channel models. At the time of analysis, 496 of the entries were implemented in the *NEURON* language (Carnevale and Hines, 2006; Hines and Carnevale, 2001), making it the most used simulation environment on the database. Customizable ion channel models are coded in *NEURON* in so-called .*mod* files (with suffix “.mod”) (Hines and Carnevale, 2000). Mod files for all *NEURON* entries were downloaded from the *ModelDB* website. Each mod file was renamed by adding the *ModelDB* ID as a prefix to the file, in the following way: *ID_name.mod*, where *ID* is the *ModelDB* ID, and *name* is the original name of the *.mod* file. 46 .*mod* files contained more than one current of the same ion type, and were separated into distinct files for each one. The suffix “icgXY” was appended to the name, where X was the ion type, and Y was the number of the current, beginning with 1 (e.g., 1234_kv_icgK2.mod for the second Kv current in file 1234_kv.mod). Furthermore, some ModelDB entries contained more than one file with the same name. These files were added separately and given unique names by appending a version number to the name - e.g. “_v2” for the second file. The total number of files collected from the database was 3495.

### Collection of metadata

Metadata information was collected using all information in journal articles and files associated with each *ModelDB* entry of interest (SOM). Each field is listed below and defined. Note that some channel models may have missing entries in the database for information that was not stated explicitly in the journal articles or *ModelDB*. Further, we stress that metadata items corresponding to the intended neuron type, brain area and animal are strictly associated to the modeling context, and are not necessarily representative of the experimental ion channels found in that particular neuron, brain area or animal.

- **ModelDB ID.** Identification number associated with each entry on *ModelDB.* All channels from the same entry have the same number.
- **PubMed ID.** PubMed citation ID of journal articles associated with this channel’s *ModelDB* entry; may contain multiple elements, and can be empty for a select few *ModelDB* entries for which no articles were found on PubMed.
- **ion type.** The ion type, or permeability, of the channel model, as listed in the journal article and .*mod* file. The following ion types were analyzed: potassium (K), sodium (Na), calcium (Ca), nonspecific (NS). If models contained more than one current, all ion types were recorded separately. Other ion types were registered but not included in the analysis.
- **gating mode/mechanism.** The dynamic simulation variable that modulates the kinetics of the model, such as voltage (v), calcium (ca), voltage and calcium (v/ca), sodium (na), chloride (cl), light (o), and g-protein coupled (g). Only v channel models were included in the analysis, with the exception of ca and v/ca models exclusively for the K ion type.
- **subtype.** The listed ion channel type, as detailed in the journal article or the *.mod* file. Subtypes were listed as mentioned without conformation to any naming convention, e.g., (Ashburner et al., 2000; Yu and Catterall, 2004). If no subtype was given, then the subtype was recorded as *not specified.* A full list of all recorded subtypes is found in Table S1.
- **author.** Listed author(s) of the *.mod* file (programmers). If authors were not specified in the *.mod* file or on *ModelDB,* we recorded the field as *not specified.*
- **animal.** The animal model (and age, if specified) emulated in simulations, either stated explicitly, or inferred from the journal article.
- **brain area/layer.** The emulated brain area and layer of the simulation, as stated explicitly, or inferred from the journal article.
- **neuron type.** The emulated neuron type of the simulation. May be several types, or listed as *general* if no neuron type was specified.
- **neuron region.** The neuron region that the ion channel is found in, divided into dendrites, soma, axon, axon hillock, or specific areas of dendrites or axon.
- **comments.** Comments from the *.mod* file itself and any other information about the channel and model from the journal article, such as previous models or experimental data that were used to constrain the model.
- **runtime [ms].** Elapsed CPU time for running 10 repetitions of a single voltage-clamp protocol (action-potential). In plots and on the web interface, we simplify model runtimes by assignment to one of four quartiles of the distribution of runtimes of all models in each class.
- **temperature.** Details about the model’s temperature dependence, and also the temperature at which simulations and/or experiments were performed as described in the journal article.
- **citations.** Estimated number of citations as available through Google Scholar, scraped monthly to update the database entries.

A total of 3495 .*mod* files were collected from *ModelDB*. 366 of these files were tools, full neuron models, or other items that do not function as ion channel models. Out of the remaining 3150 files, .*mod* files were placed into one of five groups: voltage-dependent potassium (Kv), voltage-dependent sodium (Nav), voltage-dependent calcium (Cav), calcium-dependent potassium (KCa), and hyperpolarization-activated cation (Ih). The calcium-dependent potassium group contained both ca and v/ca channel models without any distinction. These five groups accounted for 2378 files, references for which are available in the supplementary materials. Mod files that did not fit this description were omitted from the analysis. This included pumps and active dynamics (290), receptor models (370), and models with other gating dependencies or ion types (68).

We note that the groups are not only composed of channel models that are “gated” by voltage or calcium, but also more generally “dependent” upon them — we use these two notions interchangeably here. For example, inward-rectifier (IR) potassium channels are not technically gated by voltage, but they do indeed show dependence upon the voltage and so can be modeled in an analogous way. Similarly, some Ih models emulate hyperpolarization-activated cyclic nucleotide-gated (HCN) channels, but can be modeled strictly as being voltage-dependent.

### Ancestor-descendant relationships

Each channel model was linked to previous models if a relationship was listed in the journal article. We denoted the groups of models connected by these relations as “families”. This relation could be specific, along the lines of “the A channel model’s kinetics were adapted from the B channel model in a previous journal article”. Other times, the description was vague, e.g., stating that the neuron model was adapted from a previous one, with no explicit reference to ion channel kinetics. When no information was listed about previous kinetics, both in the journal article and model files themselves, the channels were assumed to have no ancestors and to have been created de novo. However, in many situations obvious similarity in .*mod* file code as verified by a `diff` command was sufficient to link models to previous ones. In these cases, relations were established even when they were not stated in the journal articles. This task was done by hand, and as such is prone to mistakes. We repeated the collection of metadata, including ancestry relations, a second time in order to correct for potential errors — we hope to correct any remaining missing or superfluous ancestral relations with the help of user submissions (ICGenealogy, 2015).

### Voltage-clamp protocol

Mod files were run individually in a *NEURON* simulation by generating a single soma compartment of length and diameter equal to 20 *μm* and cytoplasmic resistivity of 150 *Ωcm*. A passive conductance was set to 3.334 ·10^−5^*S/cm*^2^. The simulation temperature was set to 37 °C. Reversal potentials were specified separately for each ion type. Some models (172 files) featured explicit calculation of the reversal potential, so internal and external concentration values were added as extra variables to make these equivalent. Parameter values for reversal potential and ion concentrations can be found in Table S3.

A particular model was placed in the soma and a series of five voltage clamp experiments were run (Figure S3, Table S2), with the current output being recorded. Based on the desired effect of each protocol, only particular sections of the protocols were used in comparing the kinetics (Figure S3, dashed lines; also noted in Table S2). The activation protocol featured a single voltage step level, meant to capture the activation kinetics of the model. The inactivation protocol featured a varying voltage step for a long duration, followed by a second fixed voltage step, measuring the inactivation due to the first step. The deactivation protocol featured a single voltage step at a high voltage, followed by a second voltage step of varying amplitude, meant to measure the deactivation kinetics that occur as the voltage is changed from one level to the other. The ramp protocol featured a series of four up and down ramping voltages, at different slopes. Finally, the action potential protocol features voltage deflections similar to a series of action potentials in, e.g., a L2/3 pyramidal neuron of mouse somatosensory cortex. Each of the five ion type groups featured different voltage values and durations based on differences in time constants, reversal potentials, and voltage ranges at which each ion channel class is known to be active — no quantitative comparisons were made between classes. Additionally, calcium gated channel models were simulated at seven different calcium concentrations based on known concentrations (Neher and Sakaba, 2008). Values were expressed in concentration as 10^−*x*^*mM*, with *x* taking the following values: 2.0,2.5,3.0,3.5,4.0,4.5,5.0. Voltage-clamp protocols are available for download from the ICG website (ICGenealogy, 2015).

A substantial number of .*mod* files (952 files) had to be slightly modified to work with the procedure, in one of the following ways: (1) reversal potential was renamed and made a global variable to be accessed from .*hoc* file, (2) NONSPECIFIC CURRENT was changed to a USEION statement with the correct ion type (3) extra functions and/or data were included through .*inc* files, data tables or extra .*mod* files (4) max conductance was made nonzero (5) file was split into multiple files for each current present (6) POINTER variables were removed (7) internal temperature initialization was removed, and temperature dependence was set to use the global variable “celsius”. The changed files will be made available upon publication.

All ion channel models were taken from the *ModelDB* repository, as published, with small changes as noted above. This included the assumption that model parameters as chosen by the authors were set to reproduce a given, desired, dynamical behavior which matches experimental data or other constraints. We did not consider changing the internal parameters of given models — this would prevent any feasible comparison of the models, since most given models would be able to generate a large variety of behaviors under changing parameter settings (data not shown).

A small number of .*mod* files were omitted from the analysis due to problems with running the simulation protocols (44 files, Table S4). These included files that did not compile for unknown reasons, and files that produced abnormal oscillations, or extreme values. The count of these files was 16 for Kv, 20 for Nav, 5 for Cav, 2 for KCa, and 1 for Ih.

### Data extraction & processing

The most recent version at the time of writing, version 7.3, of the *NEURON* language (Carnevale and Hines, 2014) was used to run the simulation protocols. All simulations were run in Ubuntu 12.04.5 on a single core of a Intel Core i7 @ 2.67GHz with 24 Gigabytes of RAM. *NEURON* models were injected with the five different voltage clamp protocols described above and integrated at a timestep of *dt* = 5*e*^−2^*ms*. The resulting current traces were normalized by dividing by the absolute value of the maximum conductance to make the result invariant to the conductance set in the channel models. We reasoned that the maximum current depends on the number of channel models in a particular area and was thus not related to the kinetic behavior of the channels. The traces were then subsampled at a resolution of 512 data points within the regions of interest stated above. For protocols containing graded steps (activation, inactivation, deactivation) the subsampled responses across all *c* graded voltage steps were appended into one representative vector of length *L* = 512 * *c*. For calcium gated channels, we performed a similar procedure for each of the *k* calcium concentrations separately, and then appended them into one representative vector of length *L* = 512 * *k* * *c*. See Figure S3F for a schematic of this process.

### Similarity measure

To remove the time dependence of current response waveforms, we performed discrete principal component analysis (PCA)(Ramsay and Silverman, 2005) across the temporal dimension, similar to approaches in spike sorting(Lewicki, 1998). To this end, the subsampled and appended current responses for each protocol across all *N* channels in a family yielded a *NxL* dimensional data matrix, in which we normalized each column by Z-scoring: we subtracted its mean and then divided by its standard deviation. This matrix was then dimensionality reduced by PCA across the *L* temporal entries, where we chose the reduced dimensionality to capture 99% of the variability. To normalize the range of scores across conditions while keeping the covariance structure, we divided the score vector of each protocol by the standard deviation of all score entries of this protocol. These normalized scores (denoted by *condition scores*) were then combined into a final score vector. Further correlations across protocols were removed by again dimensionality-reducing by PCA (99% variance criterion) to yield a final score vector for each model. Since response traces were relatively noise-free, a high PCA dimensionality can be chosen to capture current response dimensions that are rare across the population of models. The precise value of the variance criterion in both PCA steps, although slightly changing the resulting scores, did not affect the clustering results reported above. Similarity between two channel models was then defined as the Euclidean distance of their dimensionality reduced scores.

In summary, the principal components calculated in the first step represented response curves along which the current traces were projected to yield intermediate scores. The second transformation was a linear mixing matrix, which combined these intermediate scores. Final scores had between 16 and 29 dimensions depending on the family analyzed, which additionally allowed the efficient storage of the characteristics of the thus compressed response properties in our database. The linear PCA transformations, once calculated, can be applied to additional channel models and their current responses and allow us to efficiently score new channels and easily evaluate them against all other channels in the database (ICGenealogy, 2015).

### Clustering

For clustering of channel scores, we used Ward’s minimal variance linkage (Ward, 1963) for hierarchical clustering, as implemented in the MATLAB Statistics Toolbox (R2015A, The MathWorks Inc., Natick, MA). This method can be used to produce a division of the set of all channel models into an arbitrary number of “similar” clusters, the number of which has to be constrained by internal criteria (we assumed no a-priori existence of classes in this dataset) (Halkidi et al., 2001). To this end we employed a range of internal clustering evaluation measures, which indicate the emergence of an appropriate number of clusters. Although the evaluation of these measures requires some heuristics, they have been well established and can guide the decision as to which number of clusters to choose. Concretely, these are: the Silhouette criterion (Rousseeuw, 1987), the Dunn index (Dunn, 1973), the Davies-Bouldin index (Davies and Bouldin, 1979), and the Calinski-Harabasz measure (Calinski and Harabasz, 1974), also implemented in the MATLAB Statistics Toolbox (R2015A, The MathWorks Inc., Natick, MA). For the Dunn index, the Silhouette index and the Calinksi-Harabasz measure, high values indicate mostly compact and well-separated clusters. The Davies-Bouldin index also indicates compactness and separation, however for low values. For details and reviews on these clustering indexes see e.g., Milligan and Cooper (1985) and Halkidi et al. (2001).

Values for the indexes and heuristics applied to arrive at the cluster numbers of the main text are given in Figures S4 and Figures S5. Due to the natural partition of our dataset into five conditions used to calculate the final score, we also included a measure of heterogeneity computed directly on the traces of each condition, which we termed the “Inner distance”. For a given condition, let 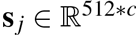 be the appended (over all possible voltage steps *c*) subsampled current responses, where *j* ∈ {1,…, *n*_channels_} runs over all channels. Let {*C_k_*|*k* ∈ {1,…, *n*_clusters_}} be a clustering of all channels — a collection of sets, such that each channel index *j* is contained in a single set. Let 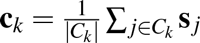 be the mean response trace of each cluster. The inner distance is then calculated as the scatter around the mean, averaged over all clusters:

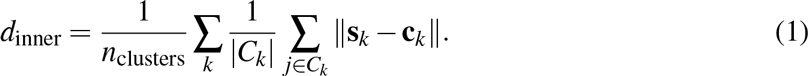

To make the measure comparable across different conditions, which might have different values of *c* (the number of voltage steps), we define the norm as 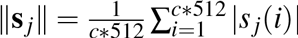.

A number of additional linkage methods (complete, single, average) and metrics (cityblock, squared euclidean) were also evaluated. While giving comparable performance on a synthetic test set, they yield mostly inferior subsections of the full set of channels with very high numbers of single elements being isolated as separate clusters.

### Assessment of protocols

To qualitatively assess the necessity of the voltage-clamp protocols for separation of labeled subtypes, the condition scores of all channel models of a particular subtype were compared with those of other subtypes (Figure S6). We show that certain protocols are more important for differentiating particular subtypes: for example, Kv models of the *m-type* show a large distinction from *A-type, dr* and *HH* subtypes in the condition scores of the action potential protocol, whereas *A-type* channel models show distinct condition scores in the activation, inactivation and deactivation protocols. The protocols chosen here thus exploit a necessary range of response kinetics; the general method of deriving a final score from each of the conditions, however, is amenable to straight-forward extension by further protocols or second-order features extracted from the response traces, as for example peak response values and time-scales (Lewicki, 1998; Druckmann et al., 2012). Each of these could be incorporated in the analysis as additional condition scores.

An alternative approach to characterizing ion channels models, given *identical implementations of models*, would be given by comparing and clustering values of the *same model parameters* across all models. Along the same lines, given a class of channel models whose kinetics all could be reproduced by a single super-model, a model-based approach would consist in fitting the parameters of this model to the responses of all ion channels models and characterizing them by the resulting values. We could not assume that the implementations of models in ModelDB are exactly the same, nor could we assume that a single super-model would capture the full dynamical diversity we were presented with, and thus we instead resorted to a “model-free” approach, as described above.

### Generation of genealogy figures

Pedigree plots were generated using Gephi (Bastian et al., 2009), and then manipulated and ordered manually for visualization (Figure 2E, Figure 4A). Coloring was chosen according to subtype label as well as cluster identity. ‘Sankey’ diagrams (Sankey, 1898; Schmidt, 2008) were generated in Javascript and D3.js (Bostock et al., 2011) (Figure 4B, Figure 5B). Subtype coloring was chosen as for the pedigree plots. Subtype labels, clusters and families were arranged from top to bottom by size. ‘Circos’ plots displaying the clustering results together with genealogical links were generated using the Circos visualization tool (Krzywinski et al., 2009), combined with TreeDyn (Chevenet et al., 2006) to create circular dendrograms (Figures 4D). Models (and clusters) were arranged on the circle in wheels using the hierarchical clustering dendrogram in its center, which places similar models (and clusters) in adjacency. Alongside the cluster and similarity relations, we plotted subtype labels, cluster majority subtypes, cluster reference models, ancestor-descendant links, runtime quartile of models and number of citations. All other figures were generated using MATLAB (R2015A, The MathWorks Inc., Natick, MA) and Python 2.7 with matplotlib 1.4.2.

### Relational database, API and web interface

All collected metadata, as well as final scores and clustering results were organized in a relational MYSQL database, which is openly queryable through a web API. Details about database structure and the implementations of the web-application and API will be made available on the website (ICGenealogy, 2015). The graphical channel browser frontend was developed in Javascript and D3.js (Bostock et al., 2011) by Phyramid Ltd, Bucharest, Romania.

### Code availability

Code for the generation of current response traces in *NEURON* as well as for the analysis of current traces will be made available on the website (ICGenealogy, 2015).

### Electrophysiology

K+ currents were recorded from Drosophila Kenyon cells in targeted in vivo whole-cell voltage clamp experiments as previously described (Murthy and Turner, 2013). Male NP7175-GAL4;UAS-mCD8-GFP flies were immobilized and fixed to a perfusion chamber using wax. Cuticle, adipose tissue, trachea and perineural sheath were removed in a window large enough to expose the posterior brain. The preparation was continuously superfused with extracellular solution containing (in mM) 103 NaCl, 3 KCl, 26 NaHCO_3_, 1 NaH_2_PO_4_, 1.5 CaCl_2_, 4 MgCl_2_, 5 TES, 8 trehalose, 10 glucose and 7 sucrose (pH 7.3 when equilibrated with 5% CO_2_ and 95% O_2_). Tetrodotoxin was added at a final concentration of 1 *μM* Borosilicate glass electrodes (14-16 MΩ) were filled with pipette solution containing (in mM) 140 potassium aspartate, 1 KCl, 10 HEPES, 4 MgATP, 0.5 Na_3_GTP and 1 EGTA (pH 7.3). All experiments were performed at room temperature (21 — 23 °C). Signals were recorded with a MultiClamp 700B Microelectrode Amplifier, lowpass-filtered at 10 and digitized at 50 kHz using a Digidata 1440A digitizer controlled via the pCLAMP 10 software (all Molecular Devices). Capacitive transients and linear leak currents were subtracted using a P/4 protocol and all traces were corrected for the liquid junction potential (Neher, 1992). Voltage pulse protocols were applied as indicated for Kv (Figure S3, Table S2) and data were analyzed in MATLAB. Resulting current traces were processed analogously to model current traces, as specified in section *Data extraction & processing*.

---

**Table S1.** Subtypes of each of the five classes currently in the database (Kv, Nav, Cav, KCa, Ih), sorted by the frequency of their occurrence. *Related to Figure 2B*.

| Kv Subtype | Count |
| --- | --- |
| A | 238 |
| delayed rectifier (dr) | 187 |
| Hodgkin-Huxley (HH) | 155 |
| m | 105 |
| not specified | 63 |
| slow | 32 |
| fast | 23 |
| inward rectifier (IR, IR2, IRK) | 17 |
| D | 15 |
| 2 | 14 |
| leak | 11 |
| 4 | 10 |
| 1 | 8 |
| RP | 7 |
| low threshold | 6 |
| high threshold | 6 |
| 3 | 5 |
| 3.1 | 5 |
| anomalous rectifier (AR) | 4 |
| x | 4 |
| 1.2 | 3 |
| persistent | 2 |
| ito | 2 |
| other | 1 |
| transient | 1 |
| 2.1 | 1 |
| KCNQ | 1 |
| 7 | 1 |
| HVA | 1 |
| udr | 1 |
| LVA | 1 |
| x1 | 1 |
| Total Models | 931 |

| Nav Subtype | Count |
| --- | --- |
| not specified | 235 |
| Hodgkin-Huxley (HH) | 115 |
| fast | 82 |
| persistent | 78 |
| resurgent | 8 |
| leak | 6 |
| 1 | 4 |
| 1.6 | 4 |
| slow | 4 |
| transient | 3 |
| 1.7 | 3 |
| 1.9 | 3 |
| 1.2 | 2 |
| T-type | 1 |
| 1.3 | 1 |
| P-type | 1 |
| 2 | 1 |
| low threshold | 1 |
| 1.8 | 1 |
| Total Models | 553 |

| Cav Subtype | Count |
| --- | --- |
| T-type | 109 |
| L-type | 99 |
| HVA | 58 |
| N-type | 33 |
| R-type | 33 |
| not specified | 25 |
| P-type | 15 |
| low threshold | 12 |
| LVA | 10 |
| high threshold | 9 |
| HH | 6 |
| TNL | 5 |
| E | 4 |
| Q | 4 |
| NPQ | 3 |
| leak | 2 |
| active flux | 1 |
| 2.1 | 1 |
| P/N | 1 |
| slow | 1 |
| Total Models | 431 |

| Ih Subtype | Count |
| --- | --- |
| h | 171 |
| HCN | 13 |
| anomalous rectifier (AR) | 7 |
| TNC | 2 |
| HH | 1 |
| Total Models | 194 |

| KCa Subtype | Count |
| --- | --- |
| not specified | 99 |
| AHP | 58 |
| BK | 34 |
| SK | 32 |
| C | 20 |
| 2 | 8 |
| CT | 6 |
| HH | 5 |
| fast | 3 |
| slow | 2 |
| mAHP | 2 |
| Total Models | 269 |

**Figure S1.**
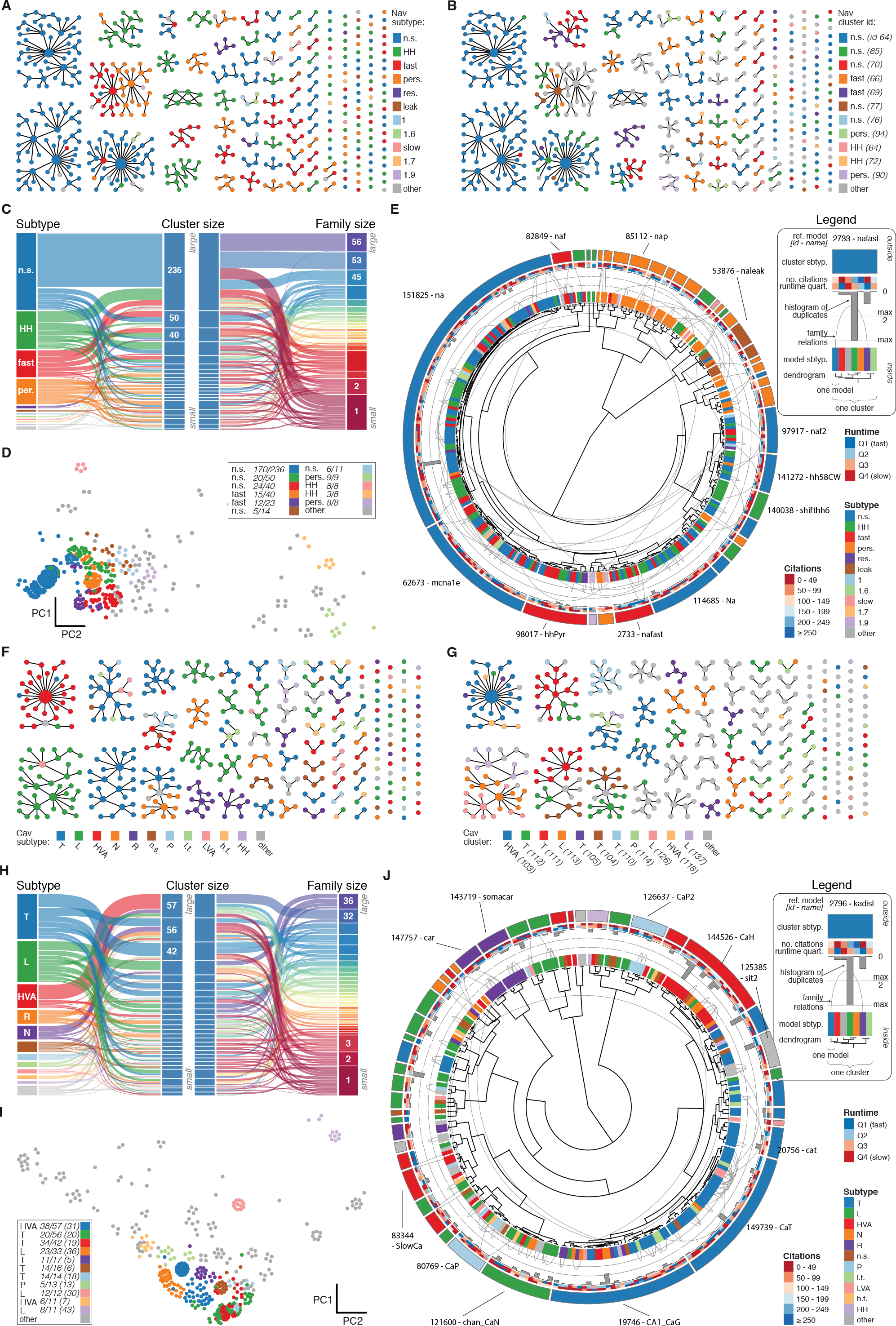
Nav and Cav class genealogy and clustering *(Related to Figure 4*). **A**: Families of the Nav class, ordered from left to right by family size. Each dot represents an ion channel model and edges represent family relationships. Colors indicate the 11 most prevalent subtypes in the class, with other subtypes colored in grey. **B** Families of the Nav class, as in panel A. Colors indicate membership in one of the 11 largest clusters in the class (named by most prevalent subtype, right hand side), with membership to other clusters colored in grey. Cluster ID is given for easy comparison with website. **C** ‘Sankey’ diagram for the Nav ion type class, showing the relation between subtype, cluster identification and family identification, each ordered from top to bottom by increasing size. The 11 most common subtypes are shown, with others being grouped together (grey). Small families (size 1 to 5 members) are grouped together. **D**: Plot of Nav models in the first two principal components of score space. Colors indicate membership in one of the 11 largest clusters in the class, with membership to other clusters colored in grey. Legend indicates the most common subtype in each cluster, and the proportion of models with that subtype. Points lying very close to each other have been distributed around the original coordinate for visualization purposes. **E**:‘Circos’ diagram of the Nav ion type class. All unique ion channel models are displayed on a ring, organized by cluster identification. From outside to inside, each segment specifies: cluster reference model (only displayed for large clusters), cluster subtype, number of citations, runtime, number of duplicates, model subtype, as well as a dendrogram of cluster connections (black) and family relations (grey). n.s.: not specified, HH: Hodgkin-Huxley, per./pers.: persistent, res.: resurgent. **F**-**J**: Cav family, similar to panels *A-E*. T: T-type, L: L-type, HVA: high-voltage activated, N: N-type, R: R-type, n.s.: not specified, P: P-type, l.t.: low threshold, LVA: low-voltage activated, h.t.: high threshold.

**Figure S2.**
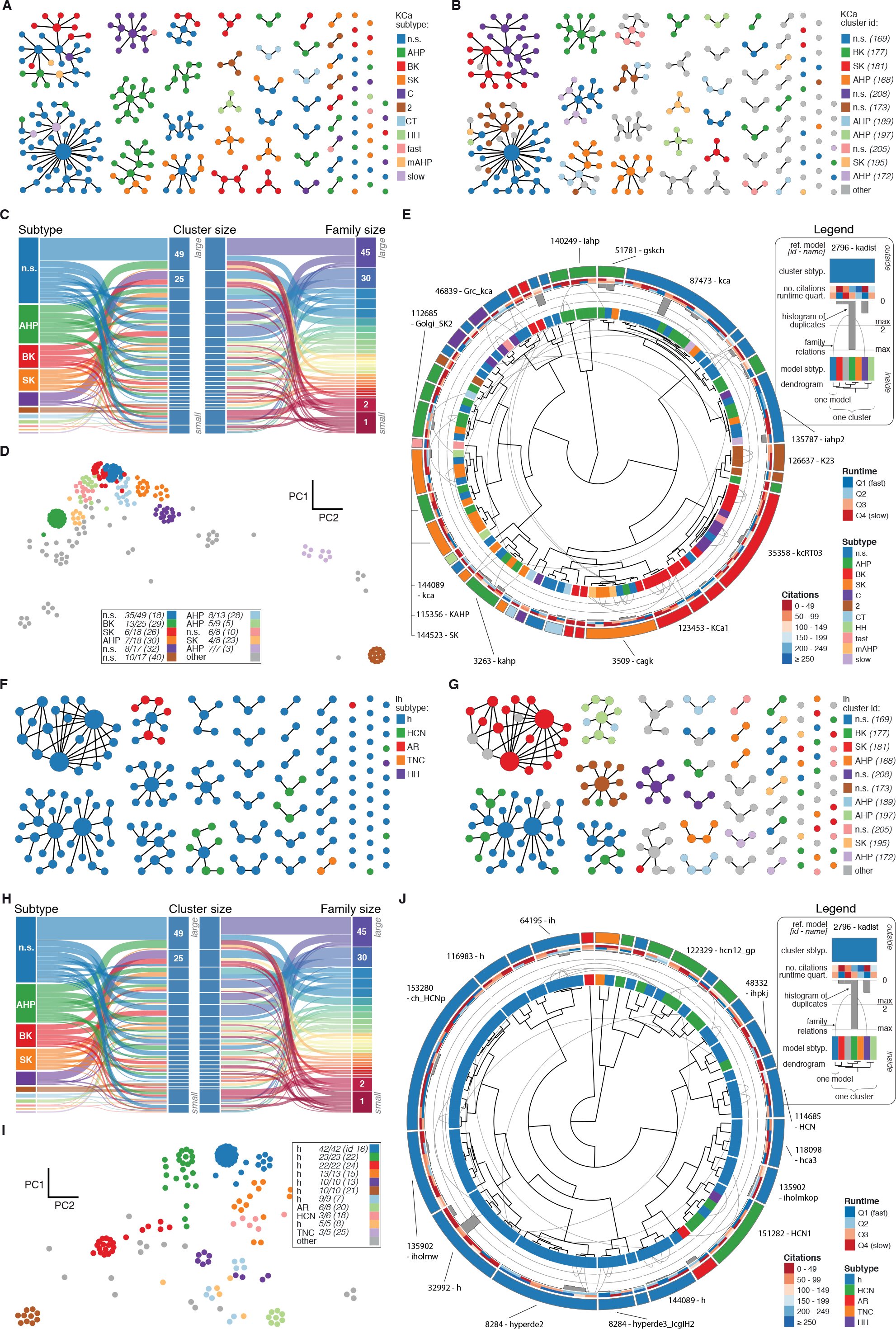
KCa and Ih class genealogy and clustering (*Related to Figure 4*). **A**: Families of the KCa class, ordered from left to right by family size. Each dot represents an ion channel model and edges represent family relationships. Colors indicate the 11 subtypes in the class. **B** Families of the KCa class, as in panel A. Colors indicate membership in one of the 11 largest clusters in the class (named by most prevalent subtype, right hand side), with membership to other clusters colored in grey. Cluster ID is given for easy comparison with website. **C** ‘Sankey’ diagram for the KCa ion type class, showing the relation between subtype, cluster identification and family identification, each ordered from top to bottom by increasing size. Small families (size 1 to 2 members) are grouped together. **D**: Plot of KCa models in the first two principal components of score space. Colors indicate membership in one of the 11 largest clusters in the class, with membership to other clusters colored in grey. Legend indicates the most common subtype in each cluster, and the proportion of models with that subtype. Points lying very close to each other have been distributed around the original coordinate for visualization purposes. **E**: ‘Circos’ diagram of the KCa ion type class. All unique ion channel models are displayed on a ring, organized by cluster identification. From outside to inside, each segment specifies: cluster reference model (only displayed for large clusters), cluster subtype, number of citations, runtime, number of duplicates, model subtype, as well as a dendrogram of cluster connections (black) and family relations (grey). n.s.: not specified, AHP: after-hyperpolarization, BK: big conductance, SK: small conductance, HH: Hodgkin-Huxley, mAHP: medium AHP. **F**-**J**: Ih family, similar to panels *A-E*. h: hyperpolarization-activated, HCN: hyperpolarization-activated cyclic nucleotide-gated, AR: anomalous rectifier, TNC: tonic nonspecific current.

**Table S2.** Voltage-clamp protocol parameters for the five ion type classes (*Related to Figure 3A*). Times are stated in units of ms, voltages in units of mV. See Figure S3 for graphical description. Items 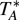 and 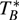 represent the starting and ending times, respectively, of the regions used for analysis (dashed areas in Figure 3, Figure S3).

| Act | Ion Type | $V_0$ | $V_1$ | $V_2$ | $\Delta V$ | $T_1$ | $T_2$ | $T_3$ | $T_A^*$ | $T_B^*$ | | | | |
| --- | --- | --- | --- | --- | --- | --- | --- | --- | --- | --- | --- | --- | --- | --- |
|  | Kv | -80 | -80 | 70 | 10 | 100 | 500 | 100 | 100 | 700 |  |  |  |  |
|  | Nav | -80 | -80 | 70 | 10 | 20 | 50 | 30 | 18 | 100 |  |  |  |  |
|  | Cav | -80 | -80 | 70 | 10 | 100 | 500 | 100 | 98 | 700 |  |  |  |  |
|  | KCa | -80 | -80 | 70 | 10 | 100 | 500 | 100 | 95 | 605 |  |  |  |  |
|  | Ih | -40 | -150 | 0 | 10 | 100 | 2000 | 100 | 95 | 2105 |  |  |  |  |
| Inact | Ion Type | $V_0$ | $V_1$ | $V_2$ | $V_3$ | $\Delta V$ | $T_1$ | $T_2$ | $T_3$ | $T_4$ | $T_A^*$ | $T_B^*$ | | |
|  | Kv | -80 | -40 | 70 | 30 | 10 | 100 | 1500 | 50 | 100 | 1600 | 1700 |  |  |
|  | Nav | -80 | -40 | 70 | 30 | 10 | 100 | 1500 | 50 | 100 | 1580 | 1750 |  |  |
|  | Cav | -80 | -40 | 70 | 30 | 10 | 100 | 1500 | 50 | 100 | 1580 | 1750 |  |  |
|  | KCa | -80 | -40 | 70 | 30 | 10 | 100 | 1500 | 50 | 100 | 1595 | 1700 |  |  |
|  | Ih | -40 | -150 | -40 | -120 | 10 | 100 | 1000 | 300 | 100 | 1095 | 1405 |  |  |
| Deact | Ion Type | $V_0$ | $V_1$ | $V_2$ | $V_3$ | $\Delta V$ | $T_1$ | $T_2$ | $T_3$ | $T_4$ | $T_A^*$ | $T_B^*$ | | |
|  | Kv | -80 | 70 | -100 | 40 | 10 | 100 | 300 | 200 | 100 | 400 | 600 |  |  |
|  | Nav | -80 | 70 | -100 | 40 | 10 | 20 | 10 | 30 | 20 | 29 | 80 |  |  |
|  | Cav | -80 | 70 | -100 | 40 | 10 | 100 | 300 | 200 | 100 | 380 | 700 |  |  |
|  | KCa | -80 | 70 | -100 | 40 | 10 | 100 | 300 | 200 | 100 | 395 | 605 |  |  |
|  | Ih | -40 | -140 | -110 | 0 | 10 | 100 | 1500 | 500 | 400 | 1595 | 2105 |  |  |
| Ramp | Ion Type | $V_0$ | $V_1$ | $T_1$ | $T_2$ | $T_3$ | $T_4$ | $T_5$ | $T_6$ | $T_7$ | $T_8$ | $T_9$ | $T_A^*$ | $T_B^*$ |
|  | Kv | -80 | 70 | 100 | 800 | 400 | 400 | 400 | 200 | 400 | 100 | 100 | 100 | 2800 |
|  | Nav | -80 | 70 | 100 | 800 | 400 | 400 | 400 | 200 | 400 | 100 | 100 | 98 | 2800 |
|  | Cav | -80 | 70 | 100 | 800 | 400 | 400 | 400 | 200 | 400 | 100 | 100 | 98 | 2800 |
|  | KCa | -80 | 70 | 100 | 800 | 400 | 400 | 400 | 200 | 400 | 100 | 100 | 100 | 2800 |
|  | Ih | -80 | 70 | 100 | 800 | 400 | 400 | 400 | 200 | 400 | 100 | 100 | 100 | 2800 |
| AP | Ion Type | $T_1$ | $T_A^*$ | $T_B^*$ | | | | | | | | | | |
|  | Kv | 1800 | 100 | 1800 |  |  |  |  |  |  |  |  |  |  |
|  | Nav | 1800 | 98 | 1800 |  |  |  |  |  |  |  |  |  |  |
|  | Cav | 1800 | 98 | 1800 |  |  |  |  |  |  |  |  |  |  |
|  | KCa | 1800 | 95 | 1655 |  |  |  |  |  |  |  |  |  |  |
|  | Ih | 1800 | 95 | 1655 |  |  |  |  |  |  |  |  |  |  |

**Table S3.** Parameters for reversal potential and inside and outside concentrations used in simulation protocols for five ion type classes (*Related to Figure 3A*). Ionic concentrations were not used for Ih currents.

| | $E_{rev}$ (mV) | $[ion]_{in}$ (mM) | $[ion]_{out}$ (mM) |
| --- | --- | --- | --- |
| Kv | -86.7 | 85.0 | 3.3152396 |
| Nav | 50.0 | 21.0 | 136.3753955 |
| Cav | 135.0 | 8.1929e-5 | 2.0 |
| KCa | -86.7 | 85.0 | 3.3152396 |
| Ih | -45.0 | - | - |

**Figure S3.**
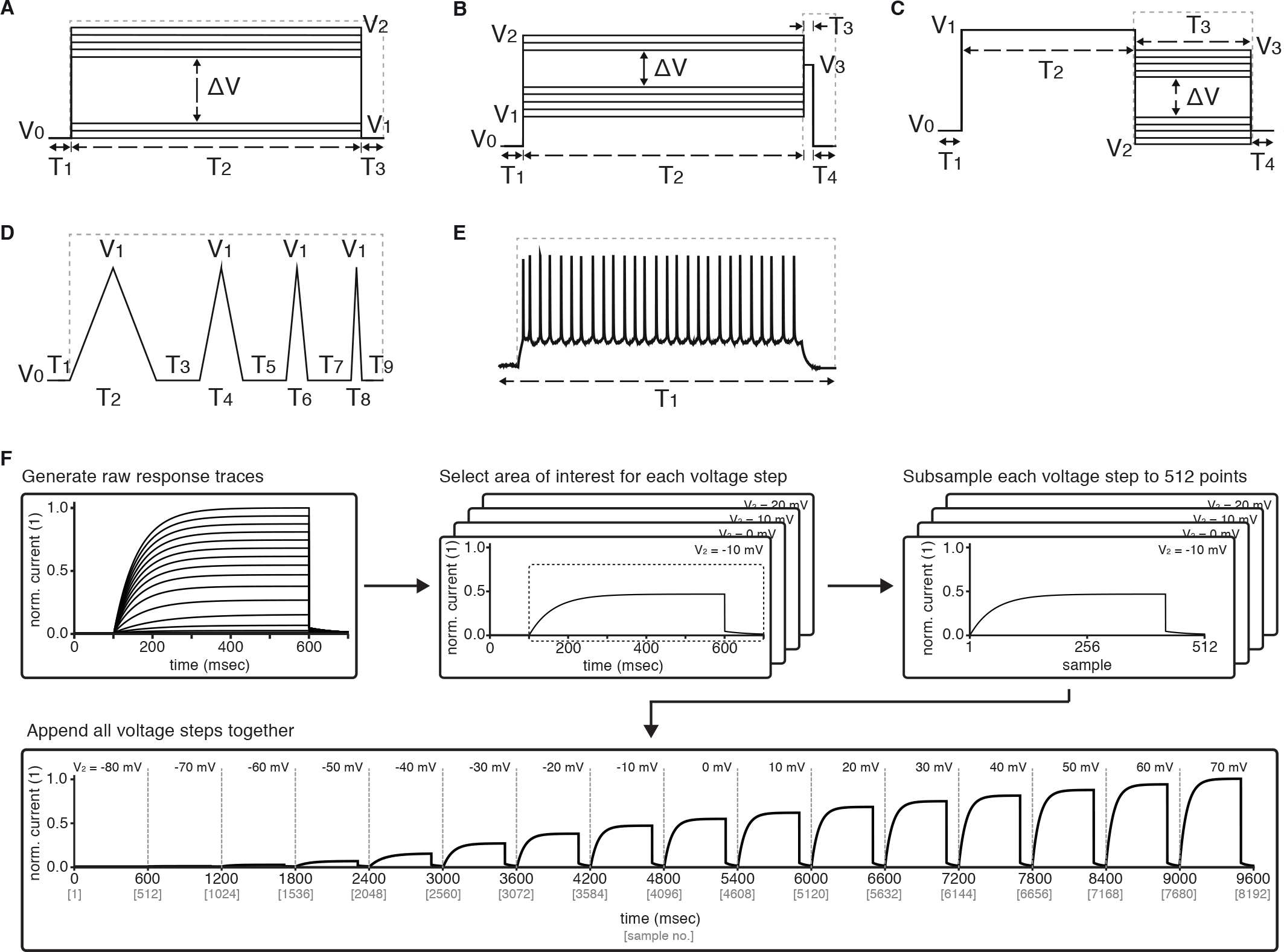
Graphical description of the five voltage-clamp protocols used for ion channel model analysis (*Related to Figure 3*): activation (**A**), inactivation (**B**), deactivation (**C**), ramp (**D**), action potential (**E**). See Table S2 for values of variables used in simulations. Dashed lines represent the approximate regions used for analysis. **F** Preprocessing stage of analysis — example shown for Kv *activation* protocol. Raw current response traces are cut to include only the region inside the dashed lines (as in panels *A-E*) for each of the (sixteen, for *activation*) voltage steps of the protocol. Then, each of the resulting traces is separately subsampled to obtain 512 data points. Finally, the subsampled traces are appended to form a final vector of size 512 * 16 = 8192. Other protocols for Kv and the other four classes are processed similarly, with the following two comments: (1) ramp and action potential protocols do not feature multiple voltage steps, and so only contain 512 datapoints total; (2) KCa current traces are run at seven different calcium concentrations, which are processed separately for each concentration as shown here, and then appended together - e.g., KCa activation will contain 512 * 16 * 7 = 57344 data points.

**Figure S4.**
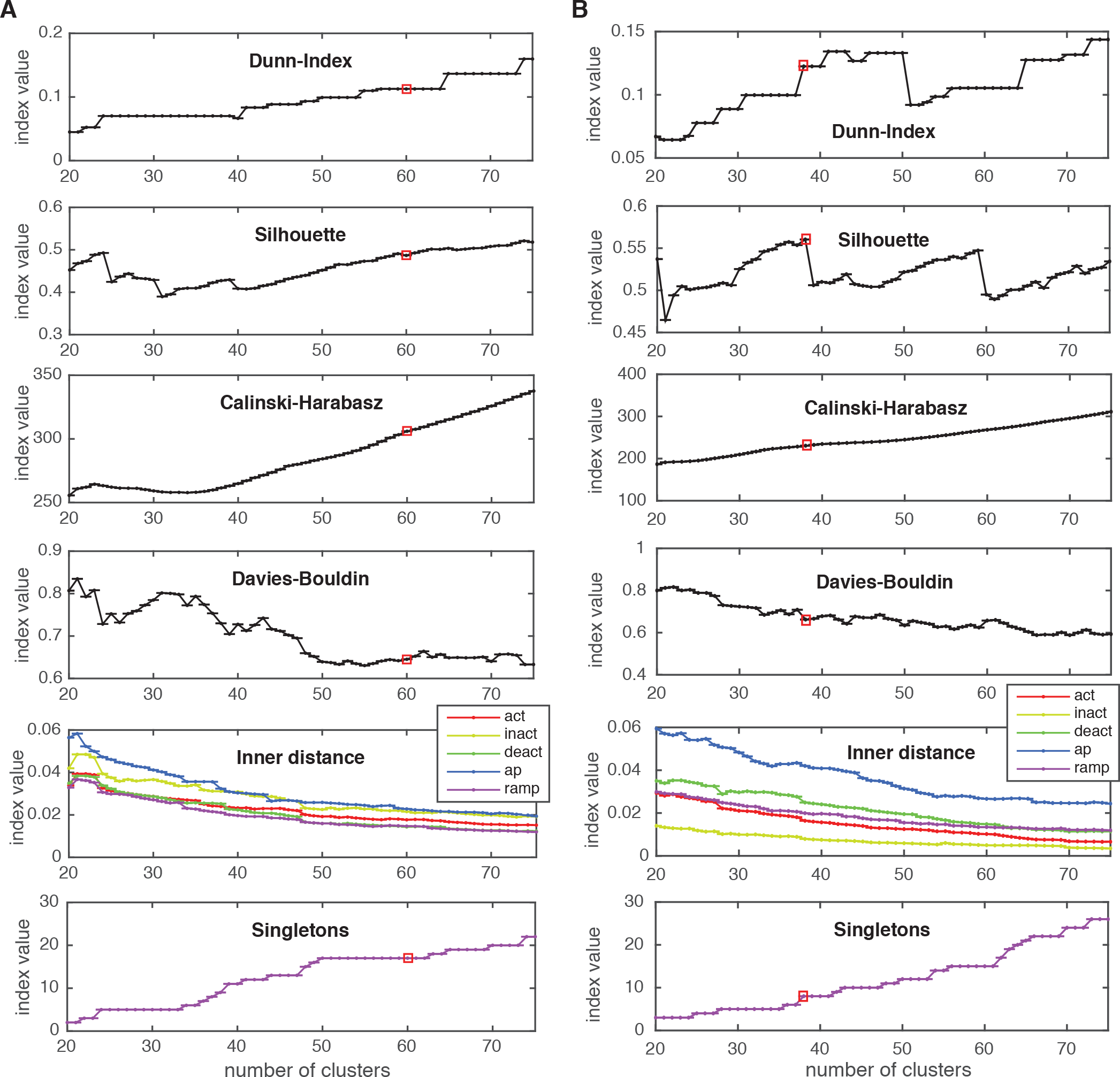
Cluster indexes, see Methods for description (*Related to Figure 3C, bottom*). “Singletons” refers to the number of single-element clusters. **A** Cluster indexes of the Kv class. The cluster number 60 was chosen since Dunn and Silhouette indexes display a large value, Calinski-Harabasz displays a “knee” (decrease in steepness) while the Davies-Bouldin is close to a local minimum. Candidate numbers 55 (Davies-Bouldin minimum) and 59 (local maximum of Silhouette, decrease in inner distance of all conditions) were disregarded due to the lower Silhouette index, and since a splitting of a major A-type cluster into two distinct clusters was observed at 60. **B** Cluster indexes of the Nav class. The cluster number 38 was chosen since Calinski-Harabasz displays a “knee” (decrease in steepness), the Dunn index displays a sharp increase, the Silhouette index a sharp decrease for higher numbers, and the inner distance displays a slight decrease in all conditions. Chosen cluster number is indicated by a red box.

**Figure S5.**
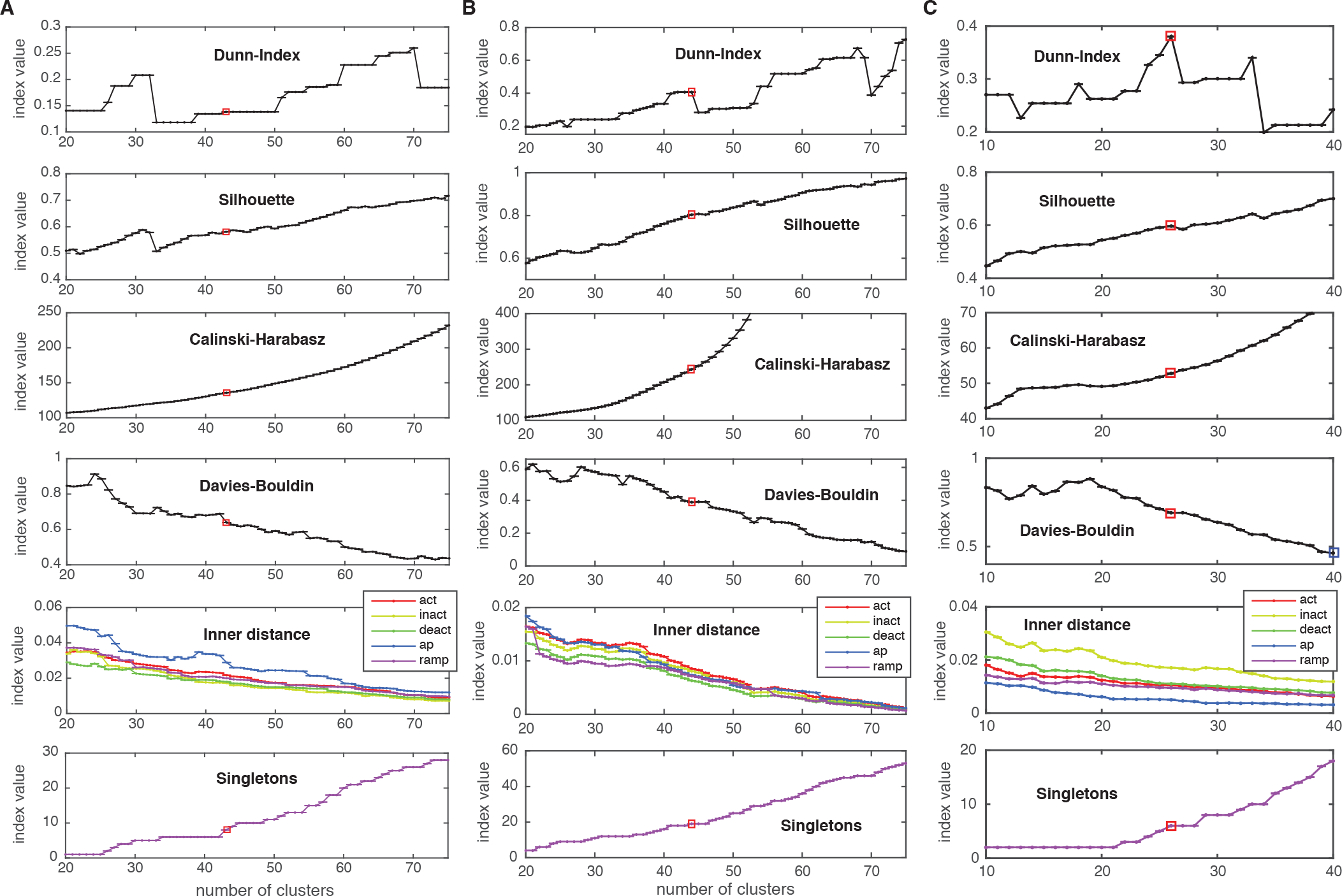
Cluster indexes, see Methods for description (*Related to Figure 3C, bottom*). “Singletons” refers to the number of single-element clusters. **A** Cluster indexes of the Cav class. The candidate cluster number 30 (peak of Dunn-index & Silhouette, local minimum of Davies-Bouldin) was disregarded since clusters were too heterogeneous (large inner distance). The next candidate cluster number 43 was chosen instead (Dunn & Silhouette display a local increase), Calinski-Harabasz displays a slight “knee” (decrease in steepness) and Davies-Bouldin displays a sharp drop. Also, a sharp decrease in “action potential” (ap) inner distance is visible at cluster number 43. **B** Cluster indexes of the KCa class. The cluster number 44 was chosen since Calinski-Harabasz displays a slight “knee” (decrease in steepness), the Dunn index displays an increase before a sharp drop while the Silhouette increases over the candidate number of 41. Also, Davies-Bouldin, stops decreasing and the inner distance displays a decrease in all conditions. **C** Cluster indexes of the Ih class. The cluster number 26 was chosen since the Dunn index displays a clear local maximum, as does the Silhouette, while Davies-Bouldin stops decreasing. The candidate number 27 (Calinski-Harabasz displays a clear “knee”) was disregarded, since the other indexes all show unfavorable changes. Chosen cluster number is indicated by a red box.

**Figure S6.**
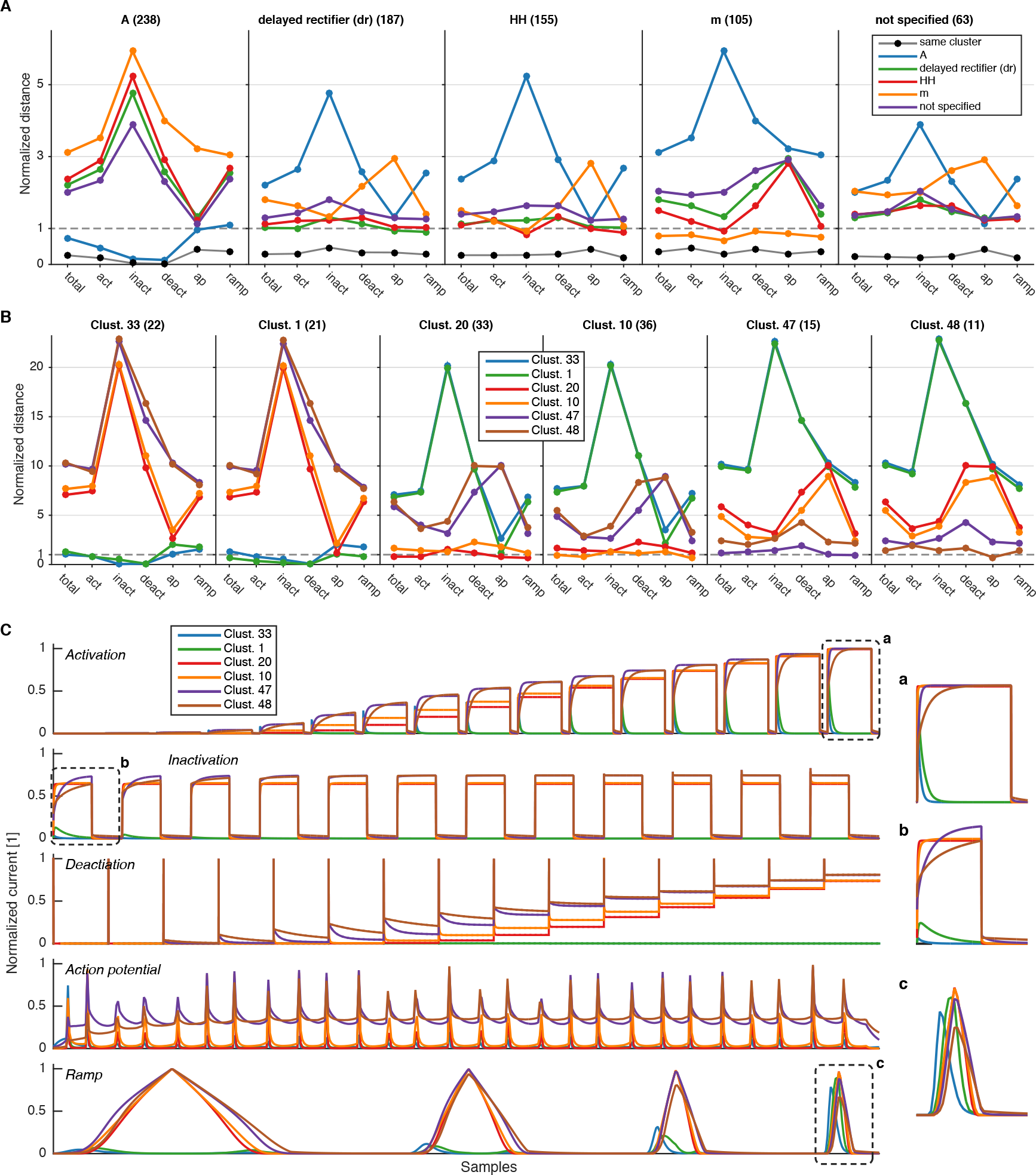
Comparison of intra– and inter-subtype variability with intra– and inter-cluster variability (*Related to Figure 3C, bottom and Figure S4A*). **A** Median pairwise distance between all channels of a given subtype, calculated either with the final score (“total”), or only each of the intermediate condition scores (here abbreviated, cf. Figure 3). Distance is given with respect to subtype given in each column, i.e. the first column displays the distance of channels of all other subtypes to the A subtype. Black circles display the median pairwise distance between those channels of a given subtype that are in the same clusters. All distances are normalized by the pairwise median distance averaged over all displayed subtypes. This shows that, e.g., *A* differs from other subtypes in all but the action potential protocol, while for example the *m-type* label differs from all other subtypes mostly in the action potential protocol. The clustering reduces the median variability to much lower levels than just the subtype partition, as visible in the black dots of each plot. Numbers in parentheses show the number of channels in the given subtype. **B** Same as in A, however pairwise distances are calculated for channels in the given clusters. Distance is given with respect to the cluster in each column, i.e. the first column displays the distance of channels in all clusters to channels in cluster 33. The first two clusters consist mostly of *A-type* channels (33: 80%,1: 91%), the middle two clusters comprise primarily of *HH/dr* channels (20: 36% HH/55% dr, 10: 39% HH/55% dr), and the right two clusters consist mostly of *m-type* channels (47: 73%,48: 73%). Numbers in parentheses correspond to the number of unique channels in the given cluster. **C** Mean current response traces of the clusters in panel B. The insets marked by a, b, c are zooms of parts of the response traces, enlarged to clarify regions of distinct current responses.

**Table S4.** List of omitted files for each of the five ion type classes, with a brief description of the reason for omission.

|  | ModelDB ID | mod name | Issue |
| --- | --- | --- | --- |
| Kv | 127992 | HHcn.mod | Produces abnormal oscillations for all protocols |
|  | 127992 | HHcnf.mod | Produces abnormal oscillations for all protocols |
|  | 141272 | hh58F1.mod | Produces abnormal current trace responses |
|  | 141272 | hh58F1ss.mod | Produces abnormal current trace responses |
|  | 155601 | markov.mod | Produces abnormal oscillations for all protocols |
|  | 155602 | markov.mod | Produces abnormal oscillations for all protocols |
|  | 135838 | GK.mod | Tab data insufficient for simulation protocol |
|  | 114637 | iA.mod | Unknown Error during run(): Segmentation Violation |
|  | 140828 | kir.mod | Table not specified in hoc func table - no table provided, not used in simulation |
| Nav | 114637 | iap.mod | Produces abnormal oscillations in AP protocol |
|  | 151443 | Naf_No.mod | Produces abnormal oscillations for all protocols |
|  | 153280 | ch_NavPVBC.mod | Produces abnormal oscillations for all protocols |
|  | 84589 | naf.mod | Produces abnormal oscillations in AP protocol |
|  | 3263 | nahh.mod | Produces abnormal oscillations for all protocols |
|  | 150446 | NafTraub1994sd.mod | Produces abnormal oscillations for all protocols |
|  | 127351 | nax.mod | Produces abnormal oscillations for all protocols |
|  | 147867 | nahh.mod | Produces abnormal oscillations for all protocols |
|  | 150239 | nax.mod | Produces abnormal oscillations for all protocols |
|  | 141272 | hh58F1.mod | Produces abnormal current trace responses |
|  | 141272 | hh58F1ss.mod | Produces abnormal current trace responses |
|  | 155601 | markov.mod | Current traces are mostly zero, does not seem to work |
|  | 155602 | markov.mod | Current traces are mostly zero, does not seem to work |
|  | 127992 | HHcn.mod | Current traces are mostly zero, does not seem to work |
|  | 127992 | HHcnf.mod | Current traces are mostly zero, does not seem to work |
|  | 135838 | GNa.mod | Tab data insufficient for simulation protocol |
| Cav | 135838 | GCa.mod | Tab data insufficient for simulation protocol |
| KCa | 139654 | kca.mod | Skipped due to mod file format (point process) |
|  | 20007 | KctBG99.mod | Produces extreme large current values, unknown reason |
| IH | 144520 | isi.mod | Unknown error during run() |

**Figure S7.**
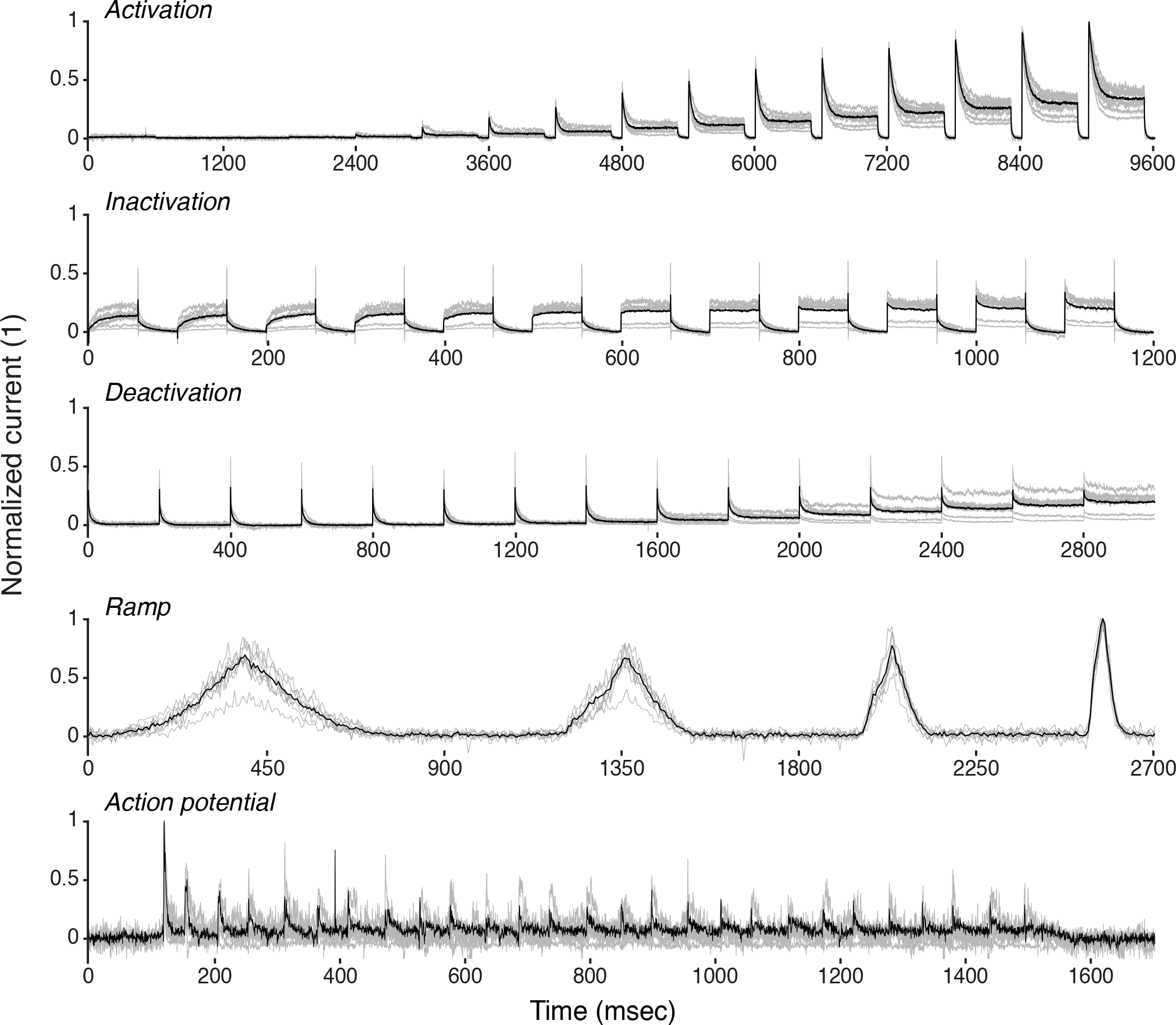
Recordings from 8 example traces of K+ current from Drosophila Kenyon cells (grey), with average (black). See Methods for a description of experiments. Response traces are arranged as detailed in Figure S3. *Related to Figure 6C*.

